# The Non-Mevalonate Pathway Requires a Delicate Balance of Intermediates to Maximize Terpene Production

**DOI:** 10.1101/2023.06.04.543646

**Authors:** Indu Raghavan, Zhen Q. Wang

## Abstract

Terpenes are valuable industrial chemicals whose demands are increasingly being met by bioengineering microbes such as *E. coli*. Although the bioengineering efforts commonly involve installing the mevalonate (MVA) pathway in *E. coli* for terpene production, the less studied methylerythritol phosphate (MEP) pathway is a more attractive target due to its higher energy efficiency and theoretical yield, despite its tight regulation. In this study, we integrated an additional copy of the entire MEP pathway into the *E. coli* genome for stable, marker-free terpene production. The genomically integrated strain produced more monoterpene geraniol than a plasmid-based system. The pathway genes’ transcription was modulated using different promoters to produce geraniol as the reporter of the pathway flux. Pathway genes, including *dxs, idi*, and *ispDF*, expressed from a medium-strength promoter, led to the highest geraniol production. Quantifying the MEP pathway intermediates revealed that the highest geraniol producers had high levels of isopentenyl pyrophosphate (IPP) and dimethylallyl pyrophosphate (DMAPP), but moderate levels of the pathway intermediates upstream of these two building blocks. A principal component analysis demonstrated that 1-deoxy-D-xylulose 5-phosphate (DXP), the product of the first enzyme of the pathway, was critical for determining the geraniol titer, whereas MEP, the product of DXP reductoisomerase (Dxr or IspC), was the least essential. This work shows that an intricate balance of the MEP pathway intermediates determines the terpene yield in engineered *E. coli*. The genetically stable and intermediate-balanced strains created in this study will serve as a chassis for producing various terpenes.

**Key Points:**

- Genome-integrated MEP pathway afforded higher strain stability
- Genome-integrated MEP pathway produced more terpene than the plasmid-based system
- High monoterpene production requires a fine balance of MEP pathway intermediates

## Introduction

Terpenes are the largest class of plant secondary metabolites with enormous commercial and medicinal values (Tetali 2019). They are often extracted from plants at low yields. Manufacturing valuable terpenes from fast-growing microbes such as *Escherichia coli* and *Saccharomyces cerevisiae* holds excellent promise for the bioeconomy (Wang et al. 2018). Two biosynthetic routes of terpenes coexist in nature: the mevalonate (MVA) pathway in eukaryotes and archaea and the methyl erythritol phosphate (MEP) pathway in bacteria and plant chloroplasts (Kuzuyama and Seto 2012). The well-studied MVA pathway is a frequent target for microbial expression to produce terpenes (Alonso-Gutierrez et al. 2013; Anthony et al. 2009; Harada et al. 2009; Kim et al. 2016; Martin et al. 2003; Rodríguez-Villalón et al. 2008; Wang et al. 2010; Wang et al. 2016; Westfall et al. 2012; Yoon et al. 2009). However, the MEP pathway is more attractive because of the higher maximum stoichiometric yield, better redox balance, and lower energy consumption (Gruchattka et al. 2013). Nevertheless, the heterologous expression of the MEP pathway in baker’s yeast is challenging due to the difficulties associated with expressing [Fe-S]-containing enzymes, in particular 4-hydroxy-3-methyl-but-2-enyl pyrophosphate synthase or IspG, in this eukaryotic host (Carlsen et al. 2013; Kirby et al. 2016; Partow et al. 2012). Thus, engineering the MEP pathway in a bacterial host such as *E. coli* is advantageous because of its ability to express functional [Fe-S]-containing enzymes.

Plasmids are frequently used to express metabolic pathway enzymes in *E. coli* due to their ease of handling. However, inserting overexpressed genes into the genome avoids metabolic burden, allows antibiotic-free cultivation, confers greater genetic stability, and maintains a precise copy number (Striedner et al. 2010; Tyo et al. 2009). Moreover, overexpressing the MEP pathway by genome integration will liberate bioengineers from repetitive engineering of the MEP pathway to increase terpene flux: the researchers can focus instead on downstream engineering by transforming plasmids encoding various prenyltransferases and terpene synthases into the genome-integrated strain to produce all kinds of terpenes. However, existing literature only reports the genomic integration of the partial MEP pathway or modulation of the promoter and RBS strengths of native MEP genes (Ajikumar et al. 2010; Li et al. 2015; Su et al. 2020; Wang et al. 2009; Yuan et al. 2006; Zhao et al. 2013a). The recent advent of the CRISPR/Cas9 mediated genome insertion in *E. coli* enables stable integration of the entire MEP pathway and the combinatorial screening of promoters to maximize terpene production.

In this study, we overexpressed the *E. coli* MEP genes by inserting an additional copy of the entire eight-gene pathway into its genome. The performance of the strains was assessed by producing a monoterpene, geraniol. The genome-integrated strain exhibited excellent genetic stability, higher cell viability, and higher geraniol titer than a strain expressing the entire pathway on plasmids. We modulated the expression of the genome-integrated pathway by creating a combinatorial library of 27 strains with promoters having different strengths. The observation that a medium-strength promoter led to the maximum geraniol titer prompted us to analyze the MEP pathway intermediates. Quantifying intracellular MEP pathway intermediates in the library revealed that not only IPP and DMAPP were crucial to terpene yield, but the delicate balance of upstream intermediates was vital to terpene yields.

## Materials and methods

### Strains, media, and culture conditions

All the strains used in this study are listed in Table S1. The strain DH5α was used for cloning all the constructs except for the final assemblies of the MEP gene cassettes, for which MG1655(DE3) was used. Chemically competent cells were transformed with plasmids using heat shock. Cells were generally plated on Luria-Bertani (LB) agar (10 g/L peptone, 5g/L yeast extract, 5 g/L NaCl, 12 g/L agar) (Thermo Fisher Scientific, Waltham, MA, USA) containing the respective antibiotics and incubated at 37°C overnight.

Single colonies were inoculated in 3 mL LB broth (10 g/L peptone, 5g/L yeast extract, and 5 g/L NaCl) (Thermo Fisher Scientific, Waltham, MA, USA) containing the respective antibiotics at 37°C at 220 rpm overnight. The overnight seed cultures were used to inoculate shake flasks containing 25 mL Terrific Broth (TB) (12 g/L tryptone, 24 g/L yeast extract, 2.2 g/L KH_2_PO_4_, and 9.4 g/L K_2_HPO_4_) (Thermo Fisher Scientific, Waltham, MA, USA) with 1.5% (w/v) glucose (MP Biomedicals, Santa Ana, CA, USA) and the respective antibiotics, at an initial OD_600_ of 0.05. The cultures were grown at 37°C at 220 rpm and induced at an OD_600_ of 1-1.2 using 1 mM isopropylthio-β-galactoside (IPTG) (GoldBio, St. Lois, MO, USA). Cultures for fluorescence imaging were grown C aerobically at 16° for 16h post-induction. The cultures for geraniol and metabolite quantification were grown at 30°C post-induction in microaerobic conditions by sealing the flasks with a parafilm. An additional 1% (w/v) glucose was added to the cultures at 24h after induction, after which flasks were re-parafilmed and grown at 30°C and harvested at various time points, as indicated in the figures.

### Molecular Cloning

All plasmids used in this study are listed in Table S2. Primers are listed in Tables S3 and S4. The ribosome binding sites for all the genes cloned in this study, except *lacI* and the antibiotic resistance genes, were designed using the RBS calculator (Salis 2011) and included in the primers. The promoters, RBSs, and terminators for *lacI* and the antibiotic resistance genes were amplified with the respective coding regions from source plasmids.

For fluorescence imaging, the genes *DsRed* and *GFPuv* were amplified from existing plasmids and ligated with the PCR-linearized pBluescript II KS+ and pCDF2 backbones, respectively, using the Gibson method (Gibson et al. 2009). *Ds*Red and GFPuv were expressed under the constitutive T7 promoter and the IPTG-inducible P_tac_ promoter, respectively. MG1655(DE3) was co-transformed with pBluescript II KS+_P_T7_-*DsRed* and pCDF2_P_tac_*-GFPuv* to create the strain with two plasmids. To construct the high copy plasmid pBluescript II KS+_P_T7_-*DsRed*.*GFPuv, DsRed* and *GFPuv* were amplified from existing plasmids and assembled as an operon under the constitutive T7 promoter into the PCR-linearized pBluescript II KS+ backbone using Gibson. The plasmid used for the genome integration of *D*sRed and *GFPuv* was created in two steps. First, pTargetF (Jiang et al. 2015) was modified to target the *araA* locus by amplifying the pTargetF backbone using 5’ phosphorylated primers containing a 20-nt sgRNA sequence homologous to the *araA* coding region to be inserted downstream of the J23119 promoter. The PCR-linearized plasmid was then circularized by the T4 ligase (NEB, Ipswich, MA, USA) to create pTargetF_*araA*. Next, the ∼500bp 5’ and 3’ homology regions of the *araA* locus were amplified from the MG1655(DE3) genome, and the operon containing *DsRed* and *GFPuv* was amplified from pBluescript II KS+_P_T7_-*DsRed*.*GFPuv*. These three gene fragments were cloned into the PCR-linearized pTargetF-*araA* using the Gibson method to create pTargetF_*araA::*P_T7_*-DsRed*.*GFPuv*. The wildtype MG1655(DE3) cells were transformed with the plasmid pTargetF_ *araA::*P_T7_-*DsRed*.*GFPuv* to create the single, medium copy plasmid system (Fig.1). Genome integration was carried out as described in the next section, “genome integration.”

**Fig.1.**
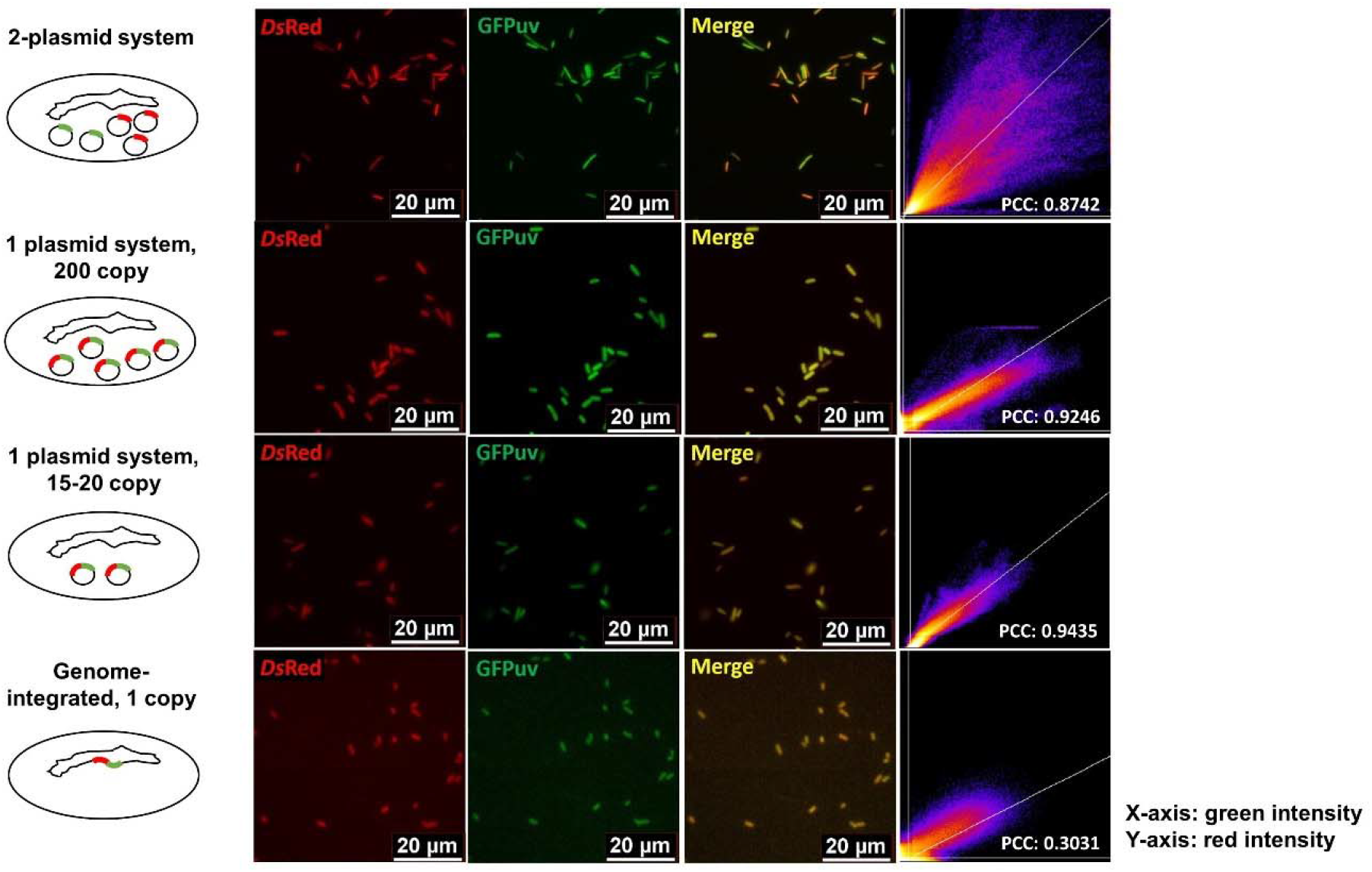
Genome-integrated expression of *Ds*Red and GFPuv showed a more significant colocalization of the two fluorescent proteins than a multi-plasmid system. Columns from left to right: schematics of different expression systems. Scatter plots of green versus red fluorescence were shown. PCC: Pearson’s Correlation Coefficient.

A modified Golden Gate method (Engler et al. 2008) was used to clone the MEP pathway genes. The plasmid pTargetF_*araA* was made Golden Gate-compatible by linearizing it using 5’ phosphorylated primers containing BsaI-HFv2 sites and then re-circularizing by ligation to create the pTargetF_*araA* GG. Plasmids pTargetF_*araA::*P_103/105/106/101/100_*-SIDF* constructed expressed *lacI*, an Anderson promoter with a lac operator, *dxs, idi, ispDF, NAT*^*R*^, and the 5’ and 3’ homology regions of the *araA* locus. All inserts were amplified using primers containing BsaI-HFv2 and Esp3I sites. Gene inserts were first assembled into the part plasmid pYTK001 using Esp3I (NEB, Ipswich, MA, USA) unless mentioned otherwise (Lee et al. 2015). Any BsaI-HFv2 and/or Esp3I sites in the inserts were domesticated prior to assembly (Mukherjee et al. 2021). *lacI* was amplified from the *E. coli* genome using primers containing the constitutive T7 promoter and BsaI-HFv2 sites and cloned into pYTK001. The IPTG-inducible Anderson and T7-lacO promoters were introduced into the vector assemblies as linear, double-stranded oligos containing BsaI-HFv2 and Esp3I sites. Genes including *dxs, idi*, and *ispDF* were amplified from the *E. coli* genome and cloned into pYTK001. The synthetic terminator, L3S2P21 (Chen et al. 2013), was linked to the last gene, *ispDF*, in the operon by including the sequence in the reverse primer. The nourseothricin-resistant gene, *NAT*^*R*^, superfolder green fluorescent protein-encoding gene (*sfGFP*), kanamycin-resistant gene (*kan*^*R*^), and the P_tac_ promoter were amplified from existing plasmids and cloned into pYTK001. The components of the integration plasmids pTargetF_*araA::*P_103/105/106/101/100_*-SIDF* were sequentially assembled into pTargetF_araA GG using BsaI-HFv2 (NEB, Ipswich, MA, USA) digestion in four steps. First, the *araA* 5’ homology region, *sfGFP*, and the *araA* 3’ homology region were assembled into pTargetF_*araA GG* to create pTargetF_*araA int GG1*. Next, the *sfGFP* was replaced by *kan*^*R*^, *ispDF*, and *NAT*^*R*^ to create pTargetF_*araA int GG2*. Next, the *kan*^*R*^ was replaced by *sfGFP, dxs*, and *idi* to create pTargetF_*araA int GG3*. Finally, the *sfGFP* was replaced by the *lacI* transcription unit and an inducible Anderson promoter to create the final plasmids pTargetF_*araA::*P_103/105/106/101/100_*-SIDF* for genome integration. All plasmids forTargetF_*pykF::*P_103/105/106/101/100/tac/T7-lacO_*-RGHE* containing the 5’ and 3’ homology regions of *pykF*, a promoter, *dxr, ispG, ispH*, and *ispE*, were cloned and assembled similarly. The native terminator, T_PheA_ (Chen et al. 2013), was placed after *ispE* by including the sequence in the primer. The reverse primers for amplifying the 5’ homology regions of both the *araA* and *pykF* loci included a stop codon to prevent the translation of any non-functional proteins. The sgRNA and homology region sequences were listed in Tables S5 and S6, respectively. The 4-nt overhang sequences used for Golden Gate assembly were listed in the SI Appendix I.

To create the plasmid pCDF2_P_tac_-*ispA(S80F)*.*tObGES* encoding the downstream genes to synthesize geraniol, a mutated *ispA* gene, *ispA(S80F)* or *ispA\**, was synthesized from Genewiz (South Plainfield, NJ, USA) and cloned into an existing plasmid containing an N-terminal truncated geraniol synthase from *Ocimum basilicum* (Iijima et al. 2004) and codon-optimized for expression in *E. coli*.

For the three-plasmid-based system for geraniol production, pTargetF_*araA::*P_106_*-SIDF* (pMB1 origin of replication, ori, spectinomycin resistance, Sp^R^) and pCDF2_P_tac_-*ispA(S80F)*.*tObGES* (200-copy *CloDF1*3 ori, Sp^R^) were modified to encode different antibiotic markers and/or origins of replication to be compatible with pTargetF_*pykF::*P_tac_*-RGHE* (pMB1 ori, Sp^R^). The antibiotic marker and origin of pTargetF_ *araA::*P_106_*-SIDF* were changed to chloramphenicol resistant gene (*cml*) and *p15A* origin, respectively, to create pIB_*araA::*P_106_*-SIDF*. To construct a plasmid expressing IspA(S80F) *and* t*Ob*GES and compatible with pIB_*araA::*P_106_*-SIDF* and pTargetF_*pykF::*P_tac_*-RGHE*, the pCDFDuet1 plasmid (*CloDF13* ori and Sp^R^) was modified to contain a kanamycin-resistant gene (*kan*^*R*^) instead of the Sp^R^ to create the pCDFDuet1-kan^R^. The PCR-amplified *ispA(S80F)* and *tObGES* were then assembled with the restriction-digested pCDFDuet1-kan^R^ (BamHI and XhoI) (NEB, Ipswich, MA, USA) using the Gibson method to create the pCDFDuet1-kan^R^_P_T7-lacO_-*ispA(S80F)*.*tObGES*. The MG1655(DE3) strain was sequentially transformed with the pIB_*araA::*P_10*6*_*-SIDF*, the pTargetF_*pykF::*P_ta*c*_*-RGHE*, and the pCDFDuet1-kan^R^_P_T7-lacO_-*ispA(S80F)*.*tObGES* to create the plasmid-based strain for geraniol production. All genes inserted into the cloned plasmids were verified by Sanger sequencing (Genewiz, South Plainfield, NJ).

### Genome integration

MG1655(DE3) was sequentially transformed with the pKD46_*Cas9*.*RecA*.*Cure* containing *SpCas9, recA*, and the genes encoding λ Red recombinases, and the pTargetF_*araA::*P_103/105/106/101/100_*-SIDF* bearing the gRNA and the MEP pathway genes to be inserted under different promoters and flanked by ∼500bp 3’ and 5’ homologous regions of *araA* locus. Cells were plated on the LB medium containing 50 μg/ml carbenicillin for pKD46_*Cas9*.*RecA*.*Cure*, 100 μg/ml spectinomycin for pTargetF_ *araA::*P_103/105/106/101/100_*-SIDF*, and 0.5% (w/v) arabinose (Acros Organics, Geel, Belgium) to induce the expression of the λ Red recombinases and grown at 30°C for 24h. Colonies were screened by PCR for the expected genome integration. The pTargetF_P_103/105/106/101/100_*-SIDF* plasmids were cured by streaking out the colony with the intended genome integration on LB plus carbenicillin but with no spectinomycin at 30°C overnight. The loss of the pTargetF_P_103/105/106/101/100_-SIDF plasmids was validated by no growth on an LB plus spectinomycin plate at 30°C overnight. The resulting strain with the *araA* integration and the pKD46_*Cas9*.*RecA*.*Cure* plasmid was then transformed with pTargetF_*pykF::*P_103/105/106/101/100/tac/T7-lacO_*-RGHE* plasmids individually and grown on an LB plate with carbenicillin, spectinomycin, and arabinose to initiate the integration at the *pykF* locus at 30°C for 24h. The resulting colonies were screened by PCR for the expected genome integration. The pTargetF_P_103/105/106/101/100/tac/T7-lacO_*-RGHE* and pKD46_*Cas9*.*RecA*.*Cure* plasmids were cured simultaneously by growing the colony with the correct integration on a plain LB plant at 37°C overnight due to the temperature-sensitive origin of replication, *repA101ts*, of pKD46_*Cas9*.*RecA*.*Cure* which degrades at 37°C. The absence of both plasmids was verified by streaking cells on LB plates containing either carbenicillin or spectinomycin.

### Fluorescence imaging

Samples for imaging were cultured at 16°C and harvested 16h after induction. Cells of OD_600_ ∼3 were centrifuged at 17,000 rpm for 2 min. The supernatants were discarded, and the pellets were resuspended in 100 μL of 0.25% (w/v) sterile melt agarose (Bio-Rad, Hercules, CA, USA). Five microliters of the samples were spotted on slides, covered with coverslips, and sealed around the edges of the coverslips using clear nail polish before imaging.

Cells were visualized on a Leica DMi 8 inverted microscope equipped with an sCMOS Leica camera (Wetzlar, Germany) using a 63x oil immersion objective. The excitation/emission wavelengths used for *Ds*Red and GFPuv were 558/583 nm and 405/510 nm, respectively. Images were individually thresholded in Fiji (ImageJ) (Schindelin et al. 2012) to minimize background fluorescence. Scatter plots were constructed using the Colocalization Finder plugin. Pearson’s Correlation Coefficients were calculated using the Just Another Colocalization Plugin (JACoP) (Bolte and Cordelières 2006).

### Gas chromatography coupled with mass spectrometry (GC/MS) for quantifying geraniol

Samples were harvested 36h post-induction. For each sample, 500 μL culture was centrifuged in a 1.5 mL microfuge tube at 17,000 rpm for 2 min. The supernatant was collected in another 1.5 mL microfuge tube to which 500 μL hexane was added. The tube was vortexed vigorously for 5 min to extract the geraniol into the hexane phase and then spun down at 17,000 rpm for 2 min. Four hundred microliters of the hexane phase containing geraniol were transferred into a glass sample vial and stored at -20°C until analysis.

A Thermo Trace 1310 Gas Chromatograph with a Thermo Scientific TraceGOLD TG-5SILMS column (30 m long, 0.25 mm inner diameter, 0.25 μm film thickness) (Thermo Fisher Scientific, Waltham, MA, USA) was used to separate the geraniol in the samples. Helium was used as the carrier gas at a flow rate of 1 mL/min. The injector was held at 200°C, and the column was injected with 5 μL of each sample using the TriPlus RSH Autosampler. The oven was initially held at 40°C for 4 min, then ramped up to 280°C at the rate of 20°C/min, and finally held at 280°C for 2 min. A Thermo Q-exactive™ Orbitrap Tandem Mass Spectrometer (Thermo Fisher Scientific, Waltham, MA, USA) was used to detect geraniol. A mass range of 39-200 m/z was monitored in the positive ion mode. Geraniol eluted at 10.24 min. Geraniol was quantified using Xcalibur™ (Thermo Fisher Scientific, Waltham, MA, USA) based on a standard curve built using 1.5 – 25 mg/L pure geraniol (Acros Organics, Geel, Belgium) in hexane. Peak areas of the ion of m/z 123.1168 ± 5 ppm were measured.

### Liquid chromatography coupled with mass spectrometry (LC/MS) for quantifying MEP pathway intermediates

Samples were harvested ∼5.5h post-induction, in the mid to late-log phase. Samples were prepared as described in a previous study (González-Cabanelas et al. 2016). Ten milliliters of culture were collected for each sample. After centrifuging at 20,000 x g for 15 min, the supernatant was discarded, and the pellet was resuspended in 1 mL methanol: acetonitrile: water (2:2:1) containing 0.1% ammonium hydroxide. After incubating on ice for 15 min and centrifuging the cells at 20,000 x g for 5 min, the supernatant containing the intracellular metabolites was extracted and dried under a stream of nitrogen in a 40°C heat block. The dried residue was dissolved in 100 μL of 10 mM ammonium acetate (adjusted to pH=9 with ammonium hydroxide) and centrifuged at 20,000 x g for 5 min. Ninety microliters of supernatant were collected in a fresh 1.5 ml microfuge tube and centrifuged for 5 min at 20,000 x g to remove any residual particles. Eighty microliters of supernatant were collected in a glass sample vial for LC/MS immediately without storage.

A Thermo UltiMate 3000 Ultra-high performance liquid chromatography system fitted with an Atlantis Premier BEH Z-HILIC VanGuard FIT column (1.7 μm particle size, 2.1 mm long, 100 mm inner diameter) (Thermo Fisher Scientific, Waltham, MA, USA) to separate the MEP pathway metabolites. The LC column compartment was maintained at 25°C. Ten microliters of each sample were injected into the column. Solution A was 50 mM ammonium carbonate in water, and solution B was acetonitrile. Isocratic elution was performed at 70% solution B. The flow rate was 0.2 mL/min. A Q-exactive Focus Orbitrap Tandem Mass Spectrometer (Thermo Fisher Scientific, Waltham, MA. USA) was used to detect the metabolites. Electrospray ionization was conducted in the negative-ion mode. The MS scan range was 50-600 m/z at a resolution of 70,000. The metabolites were quantified using Xcalibur™ based on standard curves built using 0.49 – 62.5 mg/L 1-deoxy-D-xylulose 5-phosphate (DXP), 0.49 – 125 mg/L 2-methyl-D-erythritol 4-phosphate (MEP), 0.97 – 250 mg/L 4-diphosphocytidyl-2-methylerythritol (CDP-ME), 1.95 – 1000 mg/L 2-methyl-D-erythritol-2,4-cyclopyrophosphate (MEcPP), 0.97 – 125 mg (E)-4-hydroxy-3-methyl-but-2-enyl pyrophosphate (HMBPP) (Echelon Biosciences, Inc., Salt Lake City, UT, USA) and 0.97 – 1000 mg/L dimethylallyl pyrophosphate (DMAPP) (Cayman Chemical Company, Ann Arbor, MI, USA) commercial standards prepared in 10 mM ammonium acetate were adjusted to pH=9 using ammonium hydroxide. Peak areas of the [M-H]^-^ ions of m/z 213.017 ± 5 ppm, 215.0326 ± 5 ppm, 520.0739 ± 5 ppm, 276.9884 ± 5 ppm, 260.9935 ± 5 ppm, and 244.9986 ± 5 ppm were measured for DXP, MEP, CDP-ME, MEcPP, HMBPP, and DMAPP respectively.

### Genetic stability

Geraniol-producing cell cultures were harvested 36h, 60h, and 84h post-induction. A hundred microliters of 10^4^ – 10^8^ 10-fold serial dilutions were plated on non-selective LB agar and selective LB agar containing nourseothricin for the genome-integrated strain or chloramphenicol and spectinomycin for the three-plasmid strain. After incubating at 37°C overnight, the number of colonies on each plate was counted. The colony forming unit (CFU) of each culture was calculated by multiplying the number of colonies on the respective plates with the respective dilution factors.

### Principal component analysis

Principal component analysis (PCA) was performed using packages “devtools” and “gridExtra” in R language (Chambers 2008). The metabolomic data were mean-centered and scaled to unit variance, which was calculated by subtracting each metabolite value from the mean and dividing it by the standard deviation. The complete code was listed in the “Supplemental codes” section of the supplemental information.

## Results

### Decreasing gene copy numbers resulted in uniform gene expression

Metabolic pathways are often overexpressed from one or more plasmids in *E. coli* (Liu et al. 2016; Wang et al. 2021; Yang and Guo 2014). However, the production titers are influenced by plasmid copy numbers (Ajikumar et al. 2010; Birnbaum and Bailey 1991; Silva et al. 2012). To demonstrate how plasmid-based overexpression affects pathway productivity, we created four *E. coli* strains: 1) a strain separately expressing two fluorescent proteins, *Ds*Red and GFPuv, from two different plasmids; 2) a strain expressing *DsRed* and *GFPuv* as an operon on a single high-copy plasmid (200 copies); 3) a strain expressing these two genes from one medium-copy plasmid (15-20 copies); and 4) a strain with the two genes integrated into the genome as a single operon (one copy). Fluorescence imaging showed that cells expressing *Ds*Red and GFPuv from two separate plasmids exhibited different colors, including red, green, and yellow, indicating that the genes were expressed at various levels in different cells (Fig.1). The scatter plot and Pearson’s Correlation Coefficient (PCC) showed the green and the red fluorescence varies significantly in different cells. Cells expressing the two proteins from the same plasmid or genome integration exhibited yellow colors with different intensities, indicating both proteins were expressed uniformly in each cell. The differences in color intensities reflected the differences in gene copy numbers. The lower PCC value for the genome-integrated strain was likely due to the low signal from the single copy of the genes and a high background fluorescence. Although both the single plasmid systems and the genome-integrated system resulted in a uniform expression, genome-integration was preferable for overexpressing long metabolic pathways such as the eight-gene MEP pathway because of the additional genetic stability and the limitation of cloning and transforming a large plasmid.

### Genome-integration of the complete MEP pathway

We overexpressed the complete *E. coli* MEP pathway (Fig.2a) by integrating an additional copy of the eight MEP genes into the genomic loci *araA* and *pykF*, respectively. The gene *araA* encoded L-arabinose isomerase, whose disruption prevented the breakdown of exogenous arabinose necessary to induce the expression of the lambda phage’s recombinase. The gene *pykF* encodes pyruvate kinase, whose knockout increased the MEP pathway flux by balancing pyruvate and glyceraldehyde-3-phosphate (G3P), the two precursors of the MEP pathway (Al Zaid Siddiquee et al. 2004; Farmer and Liao 2001; Li et al. 2015). The MEP pathway genes, including *dxs, idi*, and *ispDF*, were inserted into the *araA* locus (denoted as *araA*::*SIDF*), whereas *dxr, ispG, ispH*, and *ispE*, were inserted into the *pykF* locus (denoted as *pykF*::*RGHE*) as operons using the CRISPR/Cas9 guided gene editing (Fig.2b) (Jiang et al. 2015). The genome-integrated pathway was expressed under the IPTG-inducible promoters, and the lac repressor LacI was overexpressed constitutively to suppress the leaky expression from the IPTG-inducible promoters. In addition, a nourseothricin-resistant marker expressed from its native promoter was included in the *araA* locus for authentication. The engineered *E. coli* strain having the genomically integrated and overexpressed MEP pathway was transformed with a plasmid expressing IspA*, encoding an engineered geraniol pyrophosphate synthase, and t*Ob*GES, a geraniol synthase from *Ocimum basilicum* truncating the targeting peptide, to enable the synthesis of geraniol, valuable monoterpene alcohol and a precursor of the medicinally important monoterpene indole alkaloids (Brown et al. 2015; Chen and Viljoen 2010; Reiling et al. 2004).

**Fig.2.**
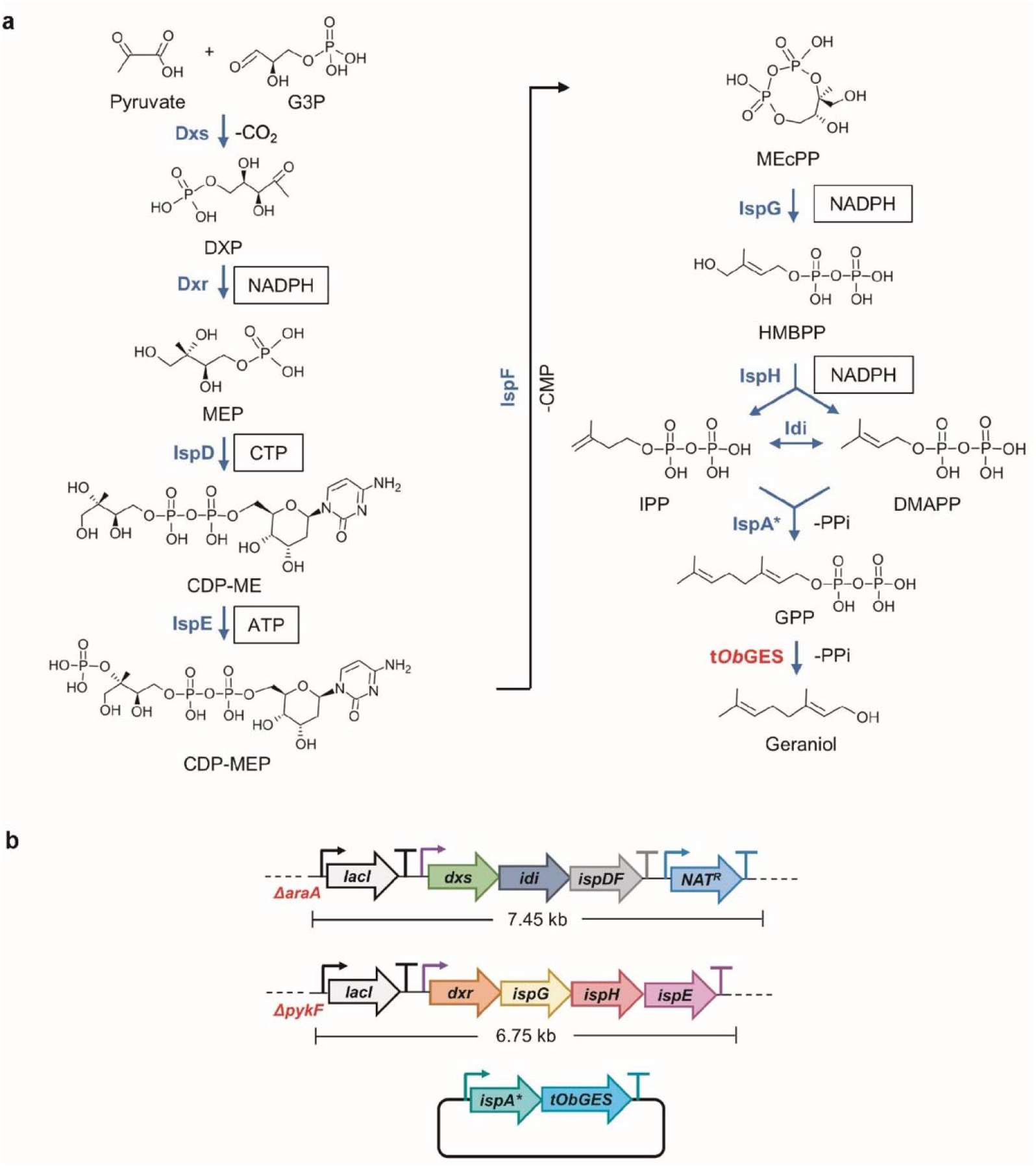
Integrating an additional copy of the MEP pathway into the *E. coli* genome for geraniol production. **a**. A schematic of the MEP pathway. The pathway enzymes were shown in blue. Dxs: 1-Deoxy-D-xylulose 5-phosphate (DXP) synthase; Dxr: 1-Deoxy-D-xylulose 5-phosphate (DXP) reductoisomerase; IspD: 2-C-Methyl-D-erythritol 4-phosphate (MEP) cytidylyltransferase, IspE: 4-Diphosphocytidyl-2-C-methylerythritol (CDP-ME) kinase, IspF: 2-C-Methyl-D-erythritol-2,4-cyclopyrophosphate (MEcPP) synthase, IspG: (E)-4-Hydroxy-3-methyl-but-2-enyl pyrophosphate (HMBPP) synthase, IspH: (E)-4-Hydroxy-3-methyl-but-2-enyl pyrophosphate (HMBPP) reductase, Idi: Isopentenyl-Diphosphate (IPP) delta isomerase 1, IspA*: *E. coli* IspA (S80F) mutant acting as a geranyl pyrophosphate synthase, t*Ob*GES: N-terminal truncated geraniol synthase from *Ocimum basilicum*. Boxed cofactors were consumed in the respective steps. NADPH: nicotinamide adenine dinucleotide phosphate, CTP: cytidine triphosphate, ATP: adenosine triphosphate, CMP: cytidine monophosphate, PPi: pyrophosphate. **b**. Schematic of the genome-integration of the MEP pathway genes. IspA* and t*Ob*GES were expressed from a plasmid. Pathway genes were expressed from various inducible Anderson promoters. LacI and Nat^R^ were expressed by constitutive promoters. IspA* and t*Ob*GES were expressed under the IPTG-inducible P_tac_ promoter. The sizes of both genome-integrated cassettes were shown.

### Genome integration afforded a higher geraniol titer compared to the plasmid-based expression

To compare the geraniol productivity from the genome-integrated system and a plasmid-based system, we created a strain overexpressing the MEP genes from two plasmids, pIB_*araA::*P_106_*-SIDF* and pTargetF_*pykF::*P_tac_*-RGHE*, respectively. The genome-integrated strain produced 5-fold more geraniol than the plasmid-based strain suggesting that the non-uniform gene expression from plasmids might hamper the product titer (Fig.3a). To determine if the genome-integrated strain was genetically more stable, we plated the liquid cultures onto solid plates with or without the antibiotics each day during the cultivation. The plasmid strain showed over 1,000 folds more colonies on the non-selective plates compared to the selective plates. This difference reflected the overwhelming loss of one or both plasmids in the plasmid-based strain, despite the antibiotics in the liquid culture. We also observed a greater loss of the pIB_*araA::*P_106_*-SIDF* plasmid compared to the pTargetF_*pykF::*P_tac_*-RGHE* plasmid, presumably due to the lower copy number of the pIB_*araA::*P_106_*-SIDF* plasmid (Fig.S1). Cells that lost plasmids died over time due to the antibiotics in the liquid culture. The cell death was evidenced by the sharply decreased cell numbers on the non-selective plate over time (Fig.3b). The genome-integrated strain was genetically stable, as expected, with similar numbers of colonies on the non-selective and selective plates at all times (Fig.3c). The cell viability of the genome-integrated strain was maintained over the course of the cultivation. The genetic instability of the plasmid-based strain was likely responsible for the decreased geraniol production compared to the genome-integrated strain.

**Fig.3.**
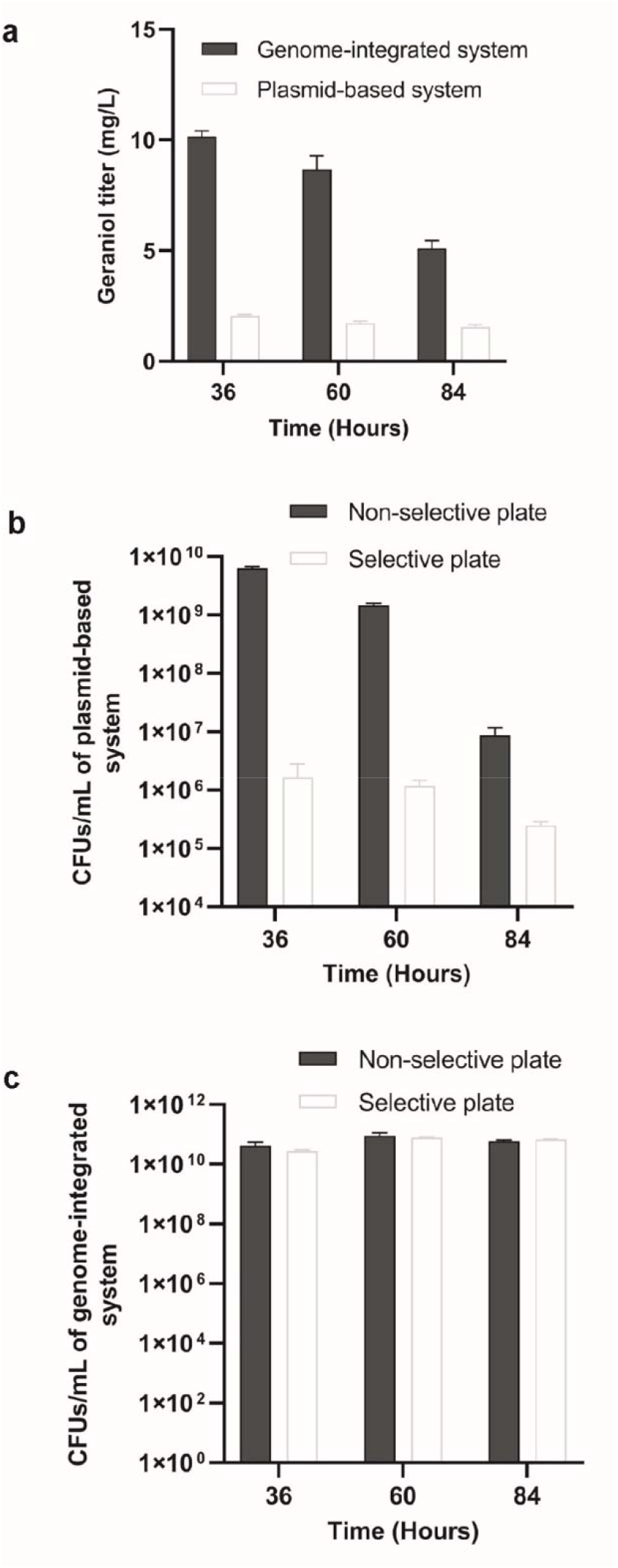
Genome-integration of the MEP pathway resulted in higher geraniol production and genetic stability. **a**. Geraniol production from the genome-integrated and the plasmid-based strains over three days. Data represent the average ± SD of three biological replicates. **b**. Number of colony forming units (CFU) per milliliter culture of the plasmid-based strain over three days. Cells were plated on non-selective plates and selective plates containing chloramphenicol and spectinomycin to select the two MEP-gene-containing plasmids. **c**. CFU per milliliter culture of the genome-integrated strain over three days. Cells were plated on non-selective plates and selective plates with nourseothricin. Cultures were diluted 10^4^∼10^8^ folds and plated in triplicates. Error bars represent the average ± S.D. of three independent biological replicates.

### Constructing a strain library to increase geraniol production and quantify MEP pathway intermediates

To optimize the MEP pathway genes’ expression, we created a 27-strain library by combinatorially testing different promoters for the integrated MEP pathway genes. IPTG-inducible Anderson promoters weaker than the P_trc_ and the T7-lacO promoters were used to avoid the metabolic burden and to titrate the promoter strengths (Balzer et al. 2013; Noh et al. 2017). Interestingly, when the promoter strength of the *araA::SIDF* increased, the geraniol titer increased first but then declined. The strains with the medium strength promoter K1585106 (P_106_) for *araA::SIDF* showed the highest geraniol production when the promoter of the *pykF::RGHE* was held constant. This observation suggested that at least one of the *araA::SIDF* enzymes, including Dxs, Idi, IspD, and IspF, was regulated beyond the transcriptional level, perhaps under feedback inhibitions. When the *araA::SIDF* was under the medium-strength P_106_ promoter, the geraniol titer increased as the promoter strength of *pykF::RGHE* increased (Fig.4a). We verified if the titer would further increase by further increasing the promoter strength of *pykF::RGHE* using strong IPTG-inducible promoters such as P_tac_ and T7-lacO. However, using the P_tac_ promoter for *pykF::RGHE* resulted in a similar level of geraniol, whereas the T7-lacO promoter decreased geraniol production, likely due to regulations downstream of transcription (Fig.4b). No obvious growth defect was observed in all engineered strains since they all had similar OD_600_ readings (Fig.S2).

**Fig.4.**
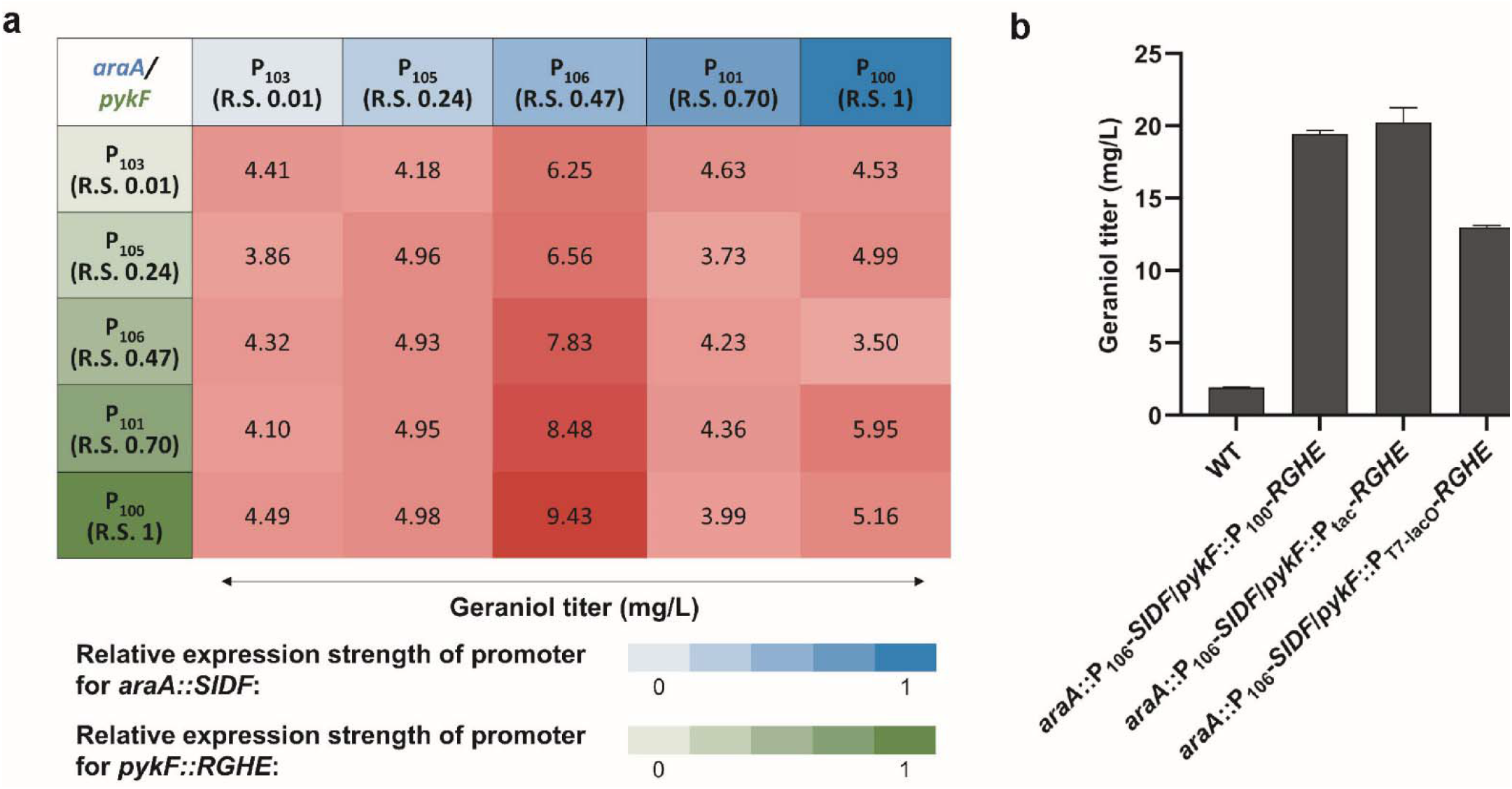
Modulating the MEP pathway gene expression to increase geraniol production. **a**. Geraniol production from a combinatorial strain library expressing the heterologous MEP pathway genes from different IPTG-inducible Anderson promoters (iGEM Registry: BBa_K1585103, BBa_K1585105, BBa_K1585106, BBa_K1585101, BBa_K1585100). Promoters expressing the *araA::SIDF* and the *pykF::RGHE* operons were shown with increasing intensities of blue and green, respectively, reflecting promoter strengths. The relative promoter strengths (R.S.) ranged from 0.01 to 1, as listed in the iGEM Registry (http://parts.igem.org/Promoters/Catalog/Anderson). Geraniol titers (mg/L) were shown with different intensities of crimson. **b**. Geraniol production from additional strains with IPTG-inducible P_tac_ and T7-lacO promoters. WT: wild type. Strain genotypes are listed in Table S1. Error bars represent the average ± S.D. of three independent biological replicates.

Because results from the combinatorial library suggested regulations beyond the transcription level, we quantified the intracellular concentrations of all MEP pathway intermediates except 4-diphosphocytidyl-2-C-methylerythritol 2-phosphate (CDP-MEP), an unstable intermediate (Li and Sharkey 2013), in 11 of the 27 strains in the combinatorial library, representing a broad range of geraniol titers. The quantification was performed during the cells’ exponential growth when the intermediate concentrations were stable and high. It was immediately noticeable that strains having high concentrations of IPP/DMAPP, the universal five-carbon terpene precursors, had medium to high geraniol titers. However, relationships between other intermediate concentrations and the geraniol titer were not obvious, thus warranting an in-depth statistical analysis.

### Principal component analysis revealed how MEP pathway intermediates correlate with geraniol production

To determine the contribution of other MEP pathway intermediates to geraniol titer, we performed a principal component analysis (PCA) using the metabolomic data gathered in Fig. 5. The strains formed three distinct clusters along the principal component 1 (PC1) and principal component 2 (PC2) axes correlating to the geraniol titers (Fig.6a). These three clusters were the top-producers, the medium-producers, and the low-producer clusters. The top producers clustered towards the lower middle part of the PCA space, while the medium producers clustered to the middle right, and the low producers were at the top left of the plot. Further analysis of the constituents of PC1 and PC2 revealed that DXP and CDP-ME were the major contributors to PC1, and MEcPP and HMBPP were the major contributors to PC2 (Fig.6b). These results demonstrated that moderate levels of DXP and CDP-ME coupled with medium-low levels of MEcPP and HMBPP were required to achieve high titers of geraniol. The geraniol production decreased with increased DXP and CDP-ME in the medium producers and further decreased with the accumulation of MEcPP and HMBPP but diminishing DXP and CDP-ME in the low producers. The wildtype strain lay at the bottom left of the PCA plot due to the low levels of all the intermediates except MEP (Fig.5 and Fig.6a). It was clear that a medium DXP level was the most critical criterion to maximize geraniol production. Additionally, MEP contributed the least to PC1 and PC2 and therefore was the least influential intermediate for geraniol production (Fig.6b). The strain *araA::*P_105_*-SIDF/pykF::*P_100_*-RGHE* was an outlier as it clustered close to the high producers but produced low levels of geraniol (Fig.5) possibly due to a technical issue during metabolite extraction. We also tried to include IPP/DMAPP with all other pathway intermediates in another PCA analysis. The resulting plot still formed three distinct clusters corresponding to the geraniol titers (Fig.S3a). Although the contributions of each intermediate were less prominent, DXP and CDP_ME were still the top two contributors to PC1, and MEcPP and HMBPP were the primary contributors to PC2 (Fig.S3b).

**Fig 5.**
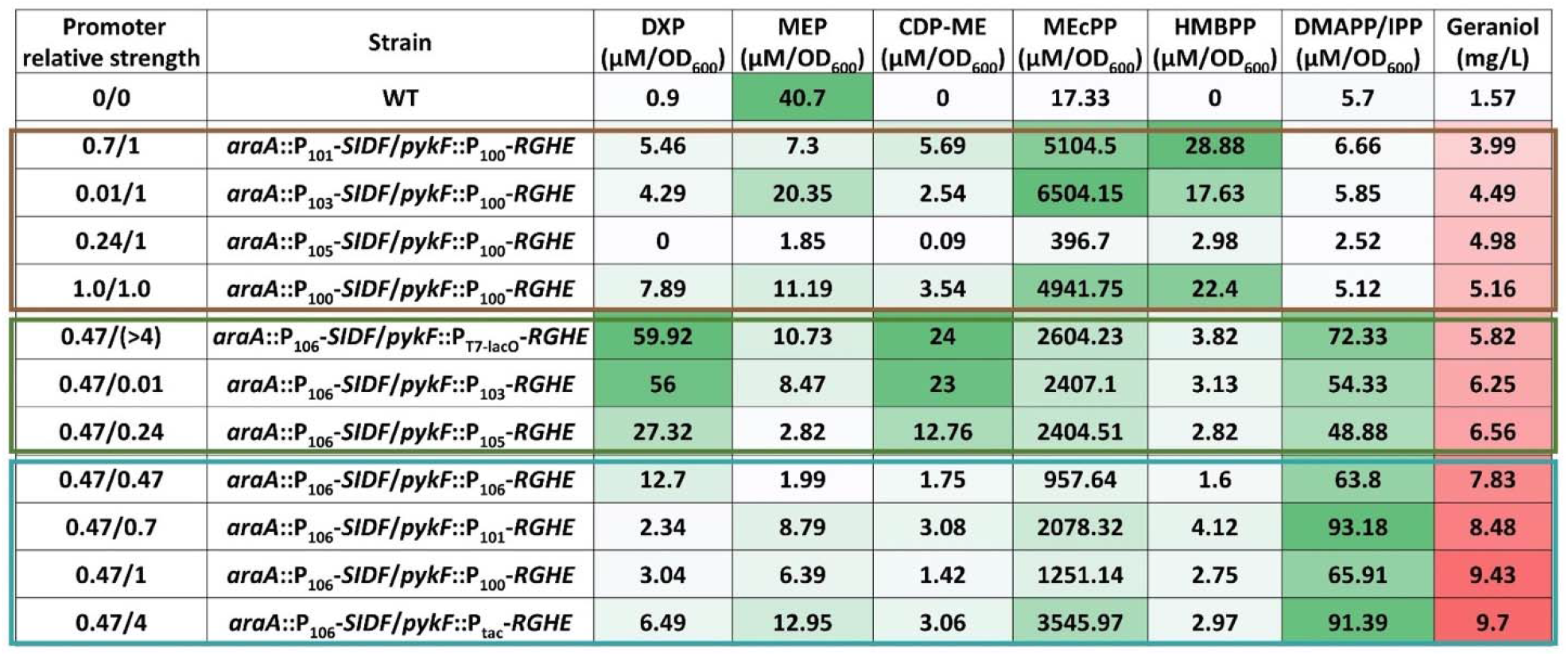
Quantifying MEP pathway intermediates in engineered *E. coli* strains. The intracellular metabolite concentrations were shown with increasing intensities in green. IPP and DMAPP were isomers that coelute. The geraniol titers were shown with increasing intensities in crimson. The numbers under “Promoter relative strength” indicated the relative strengths of promoters used for the *araA::SIDF* and the *pykF::RGHE* in each strain. The strains boxed in brown, green, and cyan were the “low producers”, “medium producers”, and “top producers”, respectively. All quantifications were performed with biological triplicates.

## Discussion

In this work, we studied the impact of overexpressing the entire MEP pathway through genome integration in *E. coli* on producing monoterpene geraniol. Though the MEP pathway has been targeted for overexpression in *E. coli* in the past, there is no report on modulating the expression of the entire genome-integrated pathway to maximize terpene titers. Integrating another copy of the MEP pathway into the *E. coli* genome has the advantage of maintaining the native MEP pathway for cell growth and the flexibility to genomically integrate heterologous MEP genes from other species. To construct a genetically stable and balanced strain for terpene production, we created a combinatorial strain library with different promoters for the genome-integrated MEP pathway and used geraniol as the reporter. Quantifying the intracellular pathway intermediate concentrations revealed the optimal levels of these intermediates to maximize the geraniol titer.

The common practice of expressing metabolic genes from multiple plasmids leads to high gene expression but also results in genetic variability and instability. Genomic integration of metabolic pathways maintains genetic stability, but the expression is limited by the gene copy number. Integrating the MVA pathway into the *E. coli* genome significantly decreased isoprenoid productivity compared to a plasmid-based expression system, partially due to the decreased gene copy number (Alonso-Gutierrez et al. 2018). Indeed, increasing the copies of genome-integrated genes by the RecA-mediated chemically induced chromosomal evolution (CIChE) produced isobutanol levels comparable to a plasmid-based expression system in *E. coli* (Saleski et al. 2021; Tyo et al. 2009). In our study, the single-copy, genome-integrated MEP pathway produced significantly more geraniol than the plasmid-based strain, which may partially attribute to the genetic variability among cells and/or dramatic plasmid loss in the plasmid-based expression system (Fig.1&3b). These observations also suggested that a fine balance of pathway intermediates, which is more achievable with genome integration than the plasmid-based system, is essential to increase the flux of a highly regulated pathway such as the MEP pathway.

The highest geraniol production in our strain library resulted from a medium strength promoter for the genes *dxs, idi*, and *ispDF* (Fig.4). This concurred with a previous study in which the highest taxadiene production resulted from relatively weak expression of the same four genes (Ajikumar et al. 2010). Indeed, the PCA analysis (Fig.6) demonstrated that medium levels of the Dxs and IspD products, DXP and CDP-ME, respectively, were required to maximize geraniol production. Dxs is the primary rate-limiting step in the MEP pathway since it showed the highest flux control coefficient (Volke et al. 2019; Wright et al. 2014). Our PCA results supported this conclusion as DXP, the product of Dxs, exerted the highest control over terpene production. Dxs, the first enzyme in this pathway, was feedback inhibited by the end products, IPP and DMAPP, typical in metabolic regulations. This allosteric regulation explained why expressing Dxs from a strong promoter decreased terpene titers (Fig.4a). A recent report delineated how IPP and DMAPP feedback inhibit Dxs: IPP and DMAPP bind to the dimerization surface of Dxs, converting the active dimer into inactive monomers, followed by protein aggregation. This elegant mechanistic elucidation enables rationally designing Dxs to circumvent this key feedback inhibition in the future (Di et al. 2022). Idi interconverts IPP and DMAPP. Although nonessential for normal bacterial growth as IspH synthesizes IPP and DMAPP simultaneously, overexpressing *idi* increased carotenoid titer and is a common target for metabolic engineering (Hahn et al. 1999). Idi can optimize the IPP: DMAPP ratio since IspH generates IPP: DMAPP at a fixed 5: 1 ratio, whereas the required IPP: DMAPP ratio varies depending on the class of terpene produced (Walsh and Tang 2017).

It is generally understood that intermediates of a pathway must be consumed effectively to prevent flux imbalances (Jones et al. 2015). Based on the metabolomic data in Fig.5, it was tempting to assume that high levels of terpene precursors, IPP and DMAPP, led to high geraniol titers. However, our PCA results showed that IPP and DMAPP levels do not solely determine geraniol production. It was essential to maintain all the intermediates upstream of IPP and DMAPP at medium or medium-low levels while maximizing the IPP and DMAPP concentrations to maximize geraniol production. The medium producers accumulated DXP and CDP-ME, preventing them from reaching high geraniol titers. Further, the low producers accumulated MEcPP and HMBPP (Fig.5 & 6a). The lack of conversion of MEcPP and HMBPP to their respective products, despite the strong overexpression of IspG and IspH in these strains, suggested that these two enzymes were limited by processes other than transcription. One of these processes could be the limited *E. coli* iron-sulfur maturation system encoded by the *isc* operon. Overexpressing the *isc* operon increased IspG and IspH’s activities by a few orders of magnitude (Gräwert et al. 2004; Zepeck et al. 2005; Zhao et al. 2013b). Thus, overexpressing the *isc* operon in the top producers will further increase terpene production in the future. The PCA analysis also identified MEP, the product of Dxr and the first committed MEP pathway intermediate, had the least influence on terpene titer. This result explained a previous observation that overexpressing Dxr did not increase carotenoid production (Yuan et al. 2006).

In conclusion, this study created an *E. coli* platform strain for terpene production by inserting an additional copy of the complete MEP pathway into the genome. The genome-integrated MEP pathway was genetically stable and reached a higher geraniol titer than a plasmid-based system. Modulating the expression of the genome-integrated MEP pathway genes revealed that a delicate balance of the pathway intermediates was critical to maximize terpene titer. In particular, IPP/DMAPP and DXP levels primarily determined final terpene yields. The highest geraniol-producing strains can be used as chassis to synthesize any terpene by transforming a plasmid containing a specific prenyltransferase and a terpene synthase.

## Acknowledgements

We are grateful to Dr. Quanjiang Ji for gifting us the plasmid pKD46_*Cas9*.*RecA*.*Cure*. We thank Dr. Valerie Frerichs for helping with the GC/MS method development and Dr. Baradwaj Ravi Gopal for helping prepare the PCA plots in figures 6 and S3.

**Fig 6.**
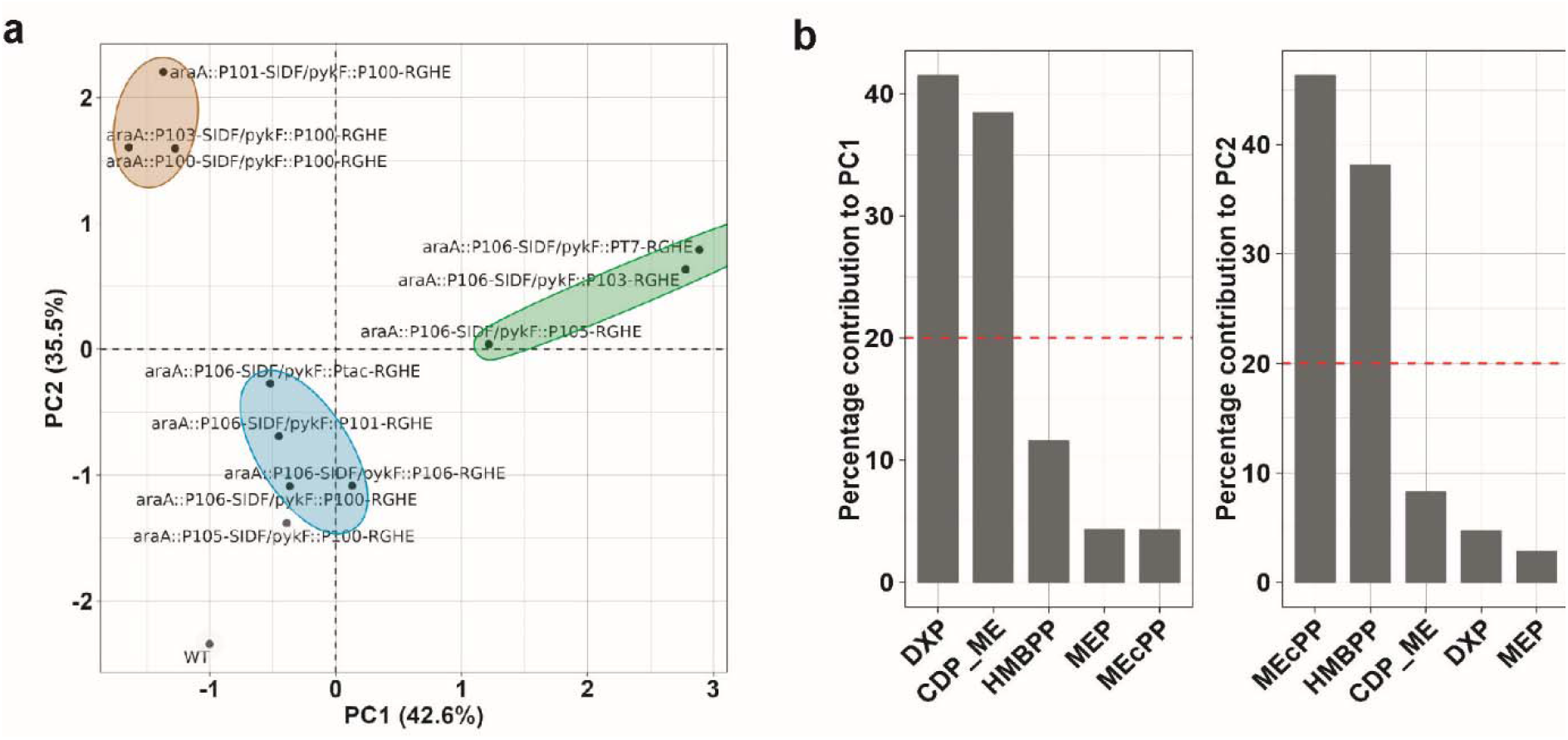
Principal component analysis (PCA) of intracellular MEP pathway intermediates except IPP/DMAPP in the engineered *E. coli* strains. **a**. Biplot of principal components (PC) 1 and 2 separating “top producers” (cyan ellipse), “medium producers” (green ellipse), and “low producers” (brown ellipse). The circled strains corresponded with the boxed strains of the same respective colors in Fig.5. WT: wild type strain. **b**. Percentage contributions of each quantified metabolite to PC1 and PC2. The dashed line indicates the expected average contribution of five variables, which is 20%, the cutoff of importance.

## Author contribution

I.R. and Z.Q.W. conceived and designed this research. I.R. conducted experiments. I.R. and Z.Q.W. analyzed data and wrote the manuscript. All authors read and approved the manuscript.

## Funding

This work was supported by the Research Foundation for the State University of New York [71272] to Z. Q. Wang, the National Science Foundation [CHE-1919594] to the University at Buffalo Chemistry Instrument Center, and the Mark Diamond Research Fund of the Graduate Student Association at the University at Buffalo, the State University of New York, to I. Raghavan.

## Data availability

All data can be provided by the corresponding author upon request.

## Statements and Declarations

### Competing Interests

The authors declare no competing interests.

### Ethical Approval

This article does not contain any studies with human participants or animals performed by any of the authors.

## Notes

### Competing Interest Statement

The authors have declared no competing interest.

## References

Ajikumar PK, Xiao W-H, Tyo KE, Wang Y, Simeon F, Leonard E, Mucha O, Phon TH, Pfeifer B, Stephanopoulos G (2010) Isoprenoid pathway optimization for Taxol precursor overproduction in Escherichia coli. Science 330(6000):70–74 doi:10.1126/science.1191652

Al Zaid Siddiquee K, Arauzo-Bravo M, Shimizu K (2004) Metabolic flux analysis of pykF gene knockout Escherichia coli based on 13 C-labeling experiments together with measurements of enzyme activities and intracellular metabolite concentrations. Appl Microbiol Biotechnol 63:407–417 doi:10.1007/s00253-003-1357-9

Alonso-Gutierrez J, Chan R, Batth TS, Adams PD, Keasling JD, Petzold CJ, Lee TS (2013) Metabolic engineering of Escherichia coli for limonene and perillyl alcohol production. Metab Eng 19:33–41 doi:10.1016/j.ymben.2013.05.004

Alonso-Gutierrez J, Koma D, Hu Q, Yang Y, Chan LJ, Petzold CJ, Adams PD, Vickers CE, Nielsen LK, Keasling JD (2018) Toward industrial production of isoprenoids in Escherichia coli: Lessons learned from CRISPR - Cas9 based optimization of a chromosomally integrated mevalonate pathway. Biotechnol Bioeng 115(4):1000–1013 doi:10.1002/bit.26530

Anthony JR, Anthony LC, Nowroozi F, Kwon G, Newman JD, Keasling JD (2009) Optimization of the mevalonate-based isoprenoid biosynthetic pathway in Escherichia coli for production of the antimalarial drug precursor amorpha-4, 11-diene. Microb Cell Factories 11(1):13–19 doi:10.1016/j.ymben.2008.07.007

Balzer S, Kucharova V, Megerle J, Lale R, Brautaset T, Valla S (2013) A comparative analysis of the properties of regulated promoter systems commonly used for recombinant gene expression in Escherichia coli. Microb Cell Factories 12(26) doi:10.1186/1475-2859-12-26

Birnbaum S, Bailey J (1991) Plasmid presence changes the relative levels of many host cell proteins and ribosome components in recombinant Escherichia coli. Biotechnol Bioeng 37(8):736–745 doi:10.1002/bit.260370808

Bolte S, Cordelières FP (2006) A guided tour into subcellular colocalization analysis in light microscopy. J Microsc 224(3):213–232 doi:10.1111/j.1365-2818.2006.01706.x

Brown S, Clastre M, Courdavault V, O’Connor SE (2015) De novo production of the plant-derived alkaloid strictosidine in yeast. Proc Natl Acad Sci USA 112(11):3205–3210 doi:10.1073/pnas.1423555112

Carlsen S, Ajikumar PK, Formenti LR, Zhou K, Phon TH, Nielsen ML, Lantz AE, Kielland-Brandt MC, Stephanopoulos G (2013) Heterologous expression and characterization of bacterial 2-C-methyl-D-erythritol-4-phosphate pathway in Saccharomyces cerevisiae. Appl Microbiol Biotechnol 97:5753–5769 doi:10.1007/s00253-013-4877-y

Chambers JM (2008) Software for data analysis: programming with R, vol 2. Springer

Chen W, Viljoen AM (2010) Geraniol—a review of a commercially important fragrance material. S Afr J Bot 76(4):643–651 doi:10.1016/j.sajb.2010.05.008

Chen Y-J, Liu P, Nielsen AA, Brophy JA, Clancy K, Peterson T, Voigt CA (2013) Characterization of 582 natural and synthetic terminators and quantification of their design constraints. Nat Methods 10(7):659–664 doi:10.1038/nmeth.2515

Di X, Ortega-Alarcon D, Kakumanu R, Iglesias-Fernandez J, Diaz L, Baidoo EE, Velazquez-Campoy A, Rodríguez-Concepción M, Perez-Gil J (2022) MEP pathway products allosterically promote monomerization of deoxy-D-xylulose-5-phosphate synthase to feedback regulate their supply. 00:100512 doi:10.1016/j.xplc.2022.100512

Engler C, Kandzia R, Marillonnet S (2008) A one pot, one step, precision cloning method with high throughput capability. PLoS One 3(11):e3647 doi:10.1371/journal.pone.0003647

Farmer WR, Liao JC (2001) Precursor balancing for metabolic engineering of lycopene production in Escherichia coli. Biotechnol Prog 17(1):57–61 doi:10.1021/bp000137t

Gibson DG, Young L, Chuang R-Y, Venter JC, Hutchison III CA, Smith HO (2009) Enzymatic assembly of DNA molecules up to several hundred kilobases. Nat Methods 6(5):343–345 doi:10.1038/nmeth.1318

González-Cabanelas D, Hammerbacher A, Raguschke B, Gershenzon J, Wright L (2016) Quantifying the metabolites of the methylerythritol 4-phosphate (MEP) pathway in plants and bacteria by liquid chromatography–triple quadrupole mass spectrometry Meth Enzymol. vol 576. Elsevier, pp 225–249

Gräwert T, Kaiser J, Zepeck F, Laupitz R, Hecht S, Amslinger S, Schramek N, Schleicher E, Weber S, Haslbeck M (2004) IspH protein of Escherichia coli: studies on Iron− Sulfur cluster implementation and catalysis. J Am Chem Soc 126(40):12847–12855 doi:10.1021/ja0471727

Gruchattka E, Hädicke O, Klamt S, Schütz V, Kayser O (2013) In silico profiling of Escherichia coli and Saccharomyces cerevisiae as terpenoid factories. Microb Cell Factories 12(84) doi:10.1186/1475-2859-12-84

Hahn FM, Hurlburt AP, Poulter CD (1999) Escherichia coli open reading frame 696 is idi, a nonessential gene encoding isopentenyl diphosphate isomerase. J Bacteriol 181(15):4499–4504 doi:10.1128/JB.181.15.4499-4504.1999

Harada H, Yu F, Okamoto S, Kuzuyama T, Utsumi R, Misawa N (2009) Efficient synthesis of functional isoprenoids from acetoacetate through metabolic pathway-engineered Escherichia coli. Appl Microbiol Biotechnol 81:915–925 doi:10.1007/s00253-008-1724-7

Iijima Y, Gang DR, Fridman E, Lewinsohn E, Pichersky E (2004) Characterization of geraniol synthase from the peltate glands of sweet basil. Plant Physiol 134(1):370–379 doi:10.1104/pp.103.032946

Jiang Y, Chen B, Duan C, Sun B, Yang J, Yang S (2015) Multigene editing in the Escherichia coli genome via the CRISPR-Cas9 system. Appl Environ Microbiol 81(7):2506–2514 doi:10.1128/AEM.04023-14

Jones JA, Toparlak ÖD, Koffas MA (2015) Metabolic pathway balancing and its role in the production of biofuels and chemicals. Curr Opin Biotechnol 33:52–59 doi:10.1016/j.copbio.2014.11.013

Kim SK, Han GH, Seong W, Kim H, Kim S-W, Lee D-H, Lee S-G (2016) CRISPR interference-guided balancing of a biosynthetic mevalonate pathway increases terpenoid production. Metab Eng 38:228–240 doi:10.1016/j.ymben.2016.08.006

Kirby J, Dietzel KL, Wichmann G, Chan R, Antipov E, Moss N, Baidoo EE, Jackson P, Gaucher SP, Gottlieb S (2016) Engineering a functional 1-deoxy-D-xylulose 5-phosphate (DXP) pathway in Saccharomyces cerevisiae. Metab Eng 38:494–503 doi:10.1016/j.ymben.2016.10.017

Kuzuyama T, Seto H (2012) Two distinct pathways for essential metabolic precursors for isoprenoid biosynthesis. Proc Jpn Acad, Ser B, Phys Biol Sci 88(3):41–52 doi:10.2183/pjab.88.41

Lee ME, DeLoache WC, Cervantes B, Dueber JE (2015) A highly characterized yeast toolkit for modular, multipart assembly. ACS Synth Biol 4(9):975–986 doi:10.1021/sb500366v

Li Y, Lin Z, Huang C, Zhang Y, Wang Z, Tang Y-j, Chen T, Zhao X (2015) Metabolic engineering of Escherichia coli using CRISPR–Cas9 meditated genome editing. Metab Eng 31:13–21 doi:10.1016/j.ymben.2015.06.006

Li Z, Sharkey TD (2013) Metabolic profiling of the methylerythritol phosphate pathway reveals the source of post-illumination isoprene burst from leaves. Plant Cell Environ 36(2):429–437 doi:10.1111/j.1365-3040.2012.02584.x

Liu W, Xu X, Zhang R, Cheng T, Cao Y, Li X, Guo J, Liu H, Xian M (2016) Engineering Escherichia coli for high-yield geraniol production with biotransformation of geranyl acetate to geraniol under fedbatch culture. Biotechnol Biofuels 9(58) doi:10.1186/s13068-016-0466-5

Martin VJ, Pitera DJ, Withers ST, Newman JD, Keasling JD (2003) Engineering a mevalonate pathway in Escherichia coli for production of terpenoids. Nat Biotechnol 21(7):796–802 doi:10.1038/nbt833

Mukherjee M, Caroll E, Wang ZQ (2021) Rapid assembly of multi-gene constructs using modular golden gate cloning. J Vis Exp 168:e61993 doi:10.3791/61993

Noh M, Yoo SM, Kim WJ, Lee SY (2017) Gene expression knockdown by modulating synthetic small RNA expression in Escherichia coli. Cell Syst 5(4):418–426. e4 doi:10.1016/j.cels.2017.08.016

Partow S, Siewers V, Daviet L, Schalk M, Nielsen J (2012) Reconstruction and evaluation of the synthetic bacterial MEP pathway in Saccharomyces cerevisiae. PLoS One 7(12):e52498 doi:10.1371/journal.pone.0052498

Reiling KK, Yoshikuni Y, Martin VJ, Newman J, Bohlmann J, Keasling JD (2004) Mono and diterpene production in Escherichia coli. Biotechnol Bioeng 87(2):200–212 doi:10.1002/bit.20128

Rodríguez-Villalón A, Pérez-Gil J, Rodríguez-Concepción M (2008) Carotenoid accumulation in bacteria with enhanced supply of isoprenoid precursors by upregulation of exogenous or endogenous pathways. J Biotechnol 135(1):78–84 doi:10.1016/j.jbiotec.2008.02.023

Saleski TE, Chung MT, Carruthers DN, Khasbaatar A, Kurabayashi K, Lin XN (2021) Optimized gene expression from bacterial chromosome by high-throughput integration and screening. Sci Adv 7(7):eabe1767 doi:10.1126/sciadv.abe1767

Salis HM (2011) The ribosome binding site calculator Meth Enzymol. vol 498. Elsevier,pp 19–42

Schindelin J, Arganda-Carreras I, Frise E, Kaynig V, Longair M, Pietzsch T, Preibisch S, Rueden C, Saalfeld S, Schmid B (2012) Fiji: an open-source platform for biological-image analysis. Nat Methods 9(7):676–682 doi:10.1038/nmeth.2019

Silva F, Queiroz JA, Domingues FC (2012) Evaluating metabolic stress and plasmid stability in plasmid DNA production by Escherichia coli. Biotechnol Adv 30(3):691–708 doi:10.1016/j.biotechadv.2011.12.005

Striedner G, Pfaffenzeller I, Markus L, Nemecek S, Grabherr R, Bayer K (2010) Plasmid-free T7-based Escherichia coli expression systems. Biotechnol Bioeng 105(4):786–794 doi:10.1002/bit.22598

Su B, Song D, Zhu H (2020) Homology-dependent recombination of large synthetic pathways into E. coli genome via λ-Red and CRISPR/Cas9 dependent selection methodology. Microb Cell Factories 19(108) doi:10.1186/s12934-020-01360-x

Tetali SD (2019) Terpenes and isoprenoids: a wealth of compounds for global use. Planta 249:1–8 doi:10.1007/s00425-018-3056-x

Tyo KE, Ajikumar PK, Stephanopoulos G (2009) Stabilized gene duplication enables long-term selection-free heterologous pathway expression. Nat Biotechnol 27(8):760–765 doi:10.1038/nbt.1555

Volke DC, Rohwer J, Fischer R, Jennewein S (2019) Investigation of the methylerythritol 4-phosphate pathway for microbial terpenoid production through metabolic control analysis. Microb Cell Factories 18(192) doi:10.1186/s12934-019-1235-5

Walsh CT, Tang Y (2017) Natural product biosynthesis. Royal Society of Chemistry

Wang C, Liwei M, Park J-B, Jeong S-H, Wei G, Wang Y, Kim S-W (2018) Microbial platform for terpenoid production: Escherichia coli and yeast. Front Microbiol 9:2460 doi:10.3389/fmicb.2018.02460

Wang C, Yoon SH, Shah AA, Chung YR, Kim JY, Choi ES, Keasling JD, Kim SW (2010) Farnesol production from Escherichia coli by harnessing the exogenous mevalonate pathway. Biotechnol Bioeng 107(3):421–429 doi:10.1002/bit.22831

Wang HH, Isaacs FJ, Carr PA, Sun ZZ, Xu G, Forest CR, Church GM (2009) Programming cells by multiplex genome engineering and accelerated evolution. Nature 460(7257):894–898 doi:10.1038/nature08187

Wang J, Niyompanich S, Tai Y-S, Wang J, Bai W, Mahida P, Gao T, Zhang K (2016) Engineering of a highly efficient Escherichia coli strain for mevalonate fermentation through chromosomal integration. Appl Environ Microbiol 82(24):7176–7184 doi:10.1128/AEM.02178-16

Wang ZQ, Song H, Koleski EJ, Hara N, Park DS, Kumar G, Min Y, Dauenhauer PJ, Chang MC (2021) A dual cellular–heterogeneous catalyst strategy for the production of olefins from glucose. Nat Chem 13(12):1178–1185 doi:10.1038/s41557-021-00820-0

Westfall PJ, Pitera DJ, Lenihan JR, Eng D, Woolard FX, Regentin R, Horning T, Tsuruta H, Melis DJ, Owens A (2012) Production of amorphadiene in yeast, and its conversion to dihydroartemisinic acid, precursor to the antimalarial agent artemisinin. Proc Natl Acad Sci USA 109(3):E111–E118 doi:10.1073/pnas.1110740109

Wright LP, Rohwer JM, Ghirardo A, Hammerbacher A, Ortiz-Alcaide M, Raguschke B, Schnitzler J-P, Gershenzon J, Phillips MA (2014) Deoxyxylulose 5-phosphate synthase controls flux through the methylerythritol 4-phosphate pathway in Arabidopsis. Plant Physiol 165(4):1488–1504 doi:10.1104/pp.114.245191

Yang J, Guo L (2014) Biosynthesis of β-carotene in engineered E. coli using the MEP and MVA pathways. Microb Cell Factories 13(160) doi:10.1186/s12934-014-0160-x

Yoon S-H, Lee S-H, Das A, Ryu H-K, Jang H-J, Kim J-Y, Oh D-K, Keasling JD, Kim S-W (2009) Combinatorial expression of bacterial whole mevalonate pathway for the production of β-carotene in E. coli. J Biotechnol 140(3-4):218–226 doi:10.1016/j.jbiotec.2009.01.008

Yuan LZ, Rouvière PE, LaRossa RA, Suh W (2006) Chromosomal promoter replacement of the isoprenoid pathway for enhancing carotenoid production in E. coli. Metab Eng 8(1):79–90 doi:10.1016/j.ymben.2005.08.005

Zepeck F, Gräwert T, Kaiser J, Schramek N, Eisenreich W, Bacher A, Rohdich F (2005) Biosynthesis of isoprenoids. Purification and properties of IspG protein from Escherichia coli. J Org Chem 70(23):9168–9174 doi:10.1021/jo0510787

Zhao J, Li Q, Sun T, Zhu X, Xu H, Tang J, Zhang X, Ma Y (2013a) Engineering central metabolic modules of Escherichia coli for improving β-carotene production. Metab Eng 17:42–50 doi:10.1016/j.ymben.2013.02.002

Zhao L, Chang W-c, Xiao Y, Liu H-w, Liu P (2013b) Methylerythritol phosphate pathway of isoprenoid biosynthesis. Annu Rev Biochem 82:497–530 doi:10.1146/annurev-biochem-052010-100934

