## Supplemental Information for "The Non-Mevalonate Pathway Requires a Delicate Balance of Intermediates to Maximize Terpene Production"

**Supplemental Methods**

*Genetic stability assay*

### The genetic stability assay in Fig.S1 was performed as follows. Cultures were grown in triplicates, and samples from each flask were harvested at 36h, 60h, and 84h post-induction. Cultures were plated on non-selective LB agar and incubated overnight at 37°C. A hundred colonies from each non-selective plate were streaked out on the respective selective media: LB+ nourseothricin for the genome-integrated strain, LB + spectinomycin, and LB + chloramphenicol for the plasmid strain to test the presence of additionally expressed MEP pathway genes. The percentage survival was calculated based on the percentage of streaks that grow on the selective plates.

**Supplemental Figures**

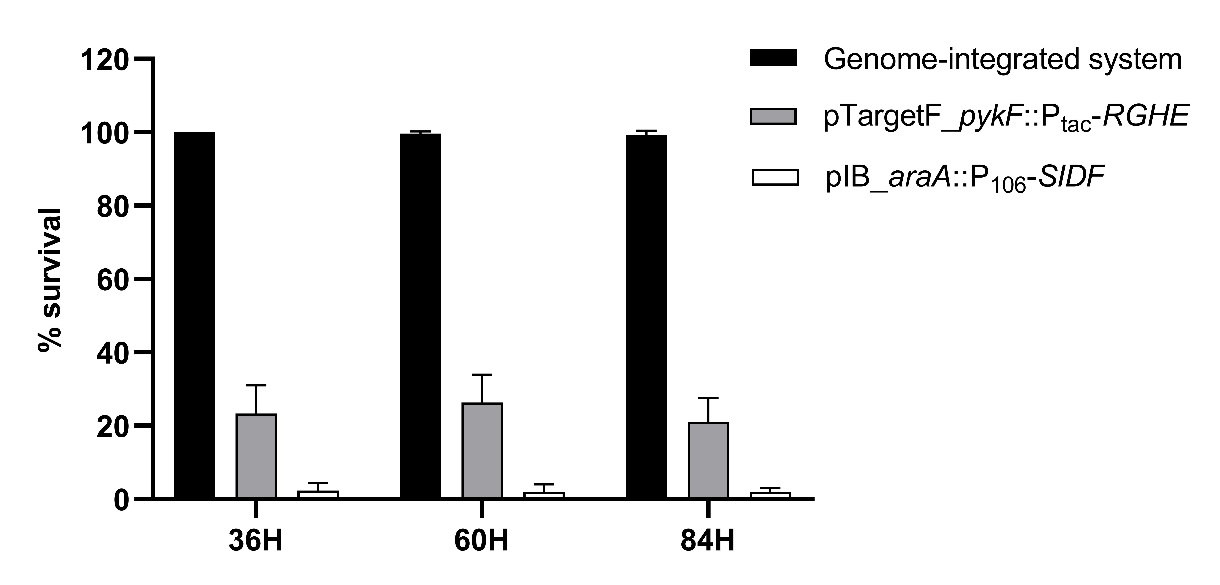

**Fig.S1** Comparison of percentage survival of strains on the respective antibiotic selective media; the genome-integrated system: Nat^R^; pIB_*araA::*P_106_-*SIDF*: Cm^R^; pTargetF_*pykF::P_tac_-RGHE*: Sp^R^. A greater loss of pIB_*araA::*P_106_-*SIDF* was seen compared to pTargetF_*pykF::P_tac_-RGHE* in the plasmid-based system. Error bars represent the average ± S.D. of three independent biological replicates.

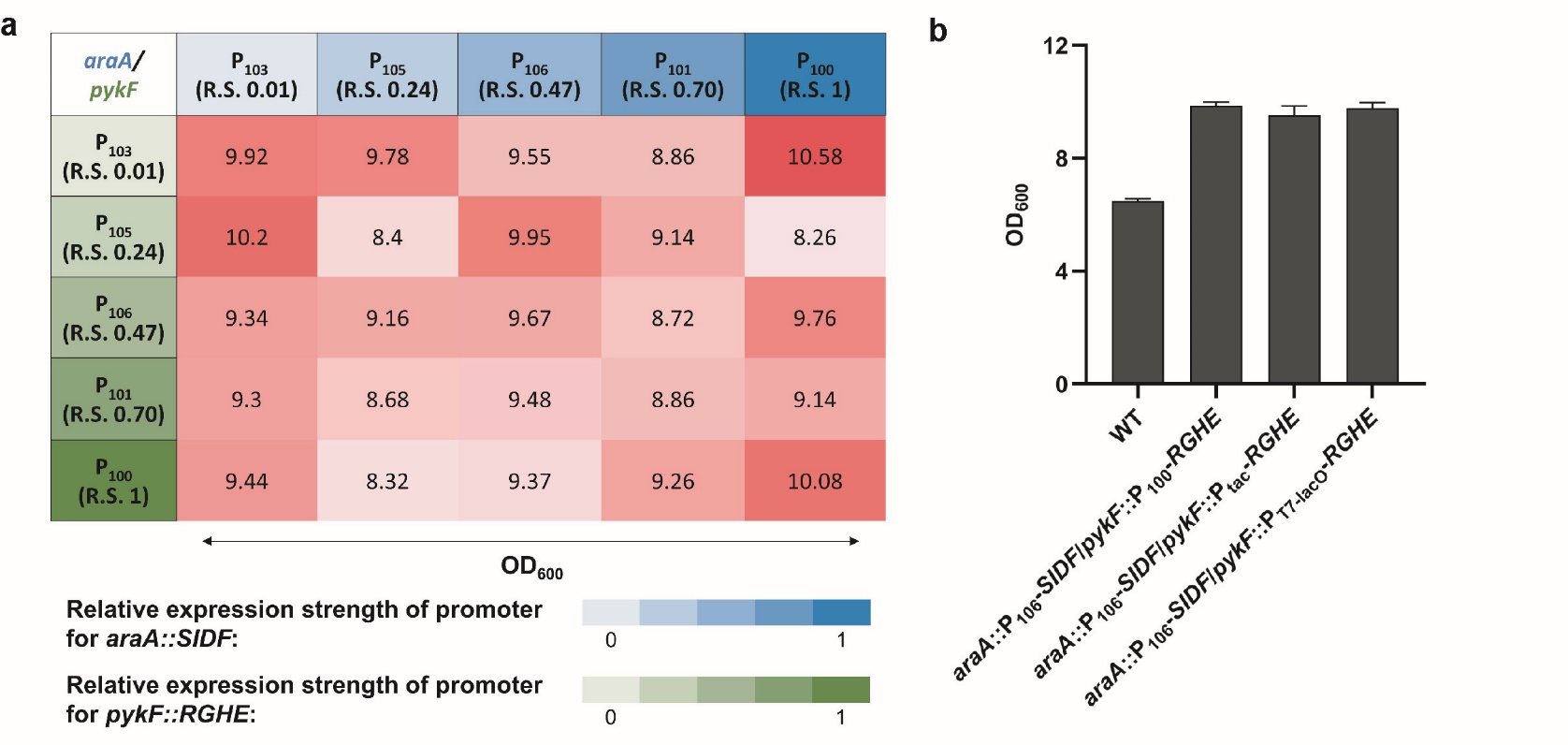

**Fig.S2** Optical densities (OD_600_) of (**a**) a combinatorial strain library expressing the heterologous MEP pathway genes from different IPTG-inducible Anderson promoters and (**b**) additional strains with IPTG-inducible P_tac_ and T7-lacO promoters. WT: wild type. Error bars represents the average ± S.D. of three independent biological replicates.

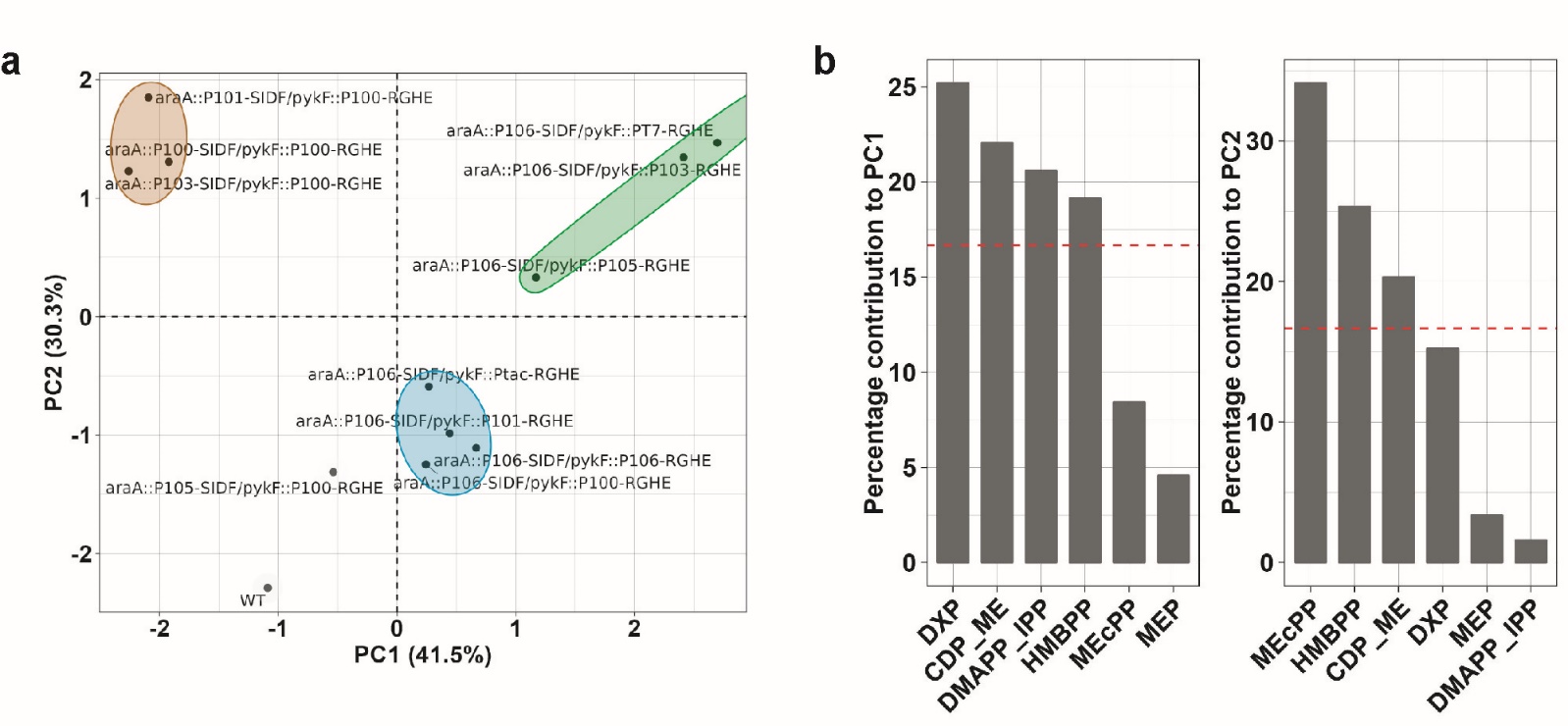

**Fig.S3** Principal component analysis (PCA) of intracellular DXP, MEP, CDP-ME, MEcPP, HMBPP, and IPP/DMAPP levels in the engineered *E. coli* strains. **a.** Biplot of principal components (PC) 1 and 2 separating “top producers” (cyan ellipse), “medium producers” (green ellipse), and “low producers” (brown ellipse). WT: Wild type. **b.** Percentage contribution of each quantified metabolite to PC1 and PC2. The dashed line indicates the expected average contribution of six variables, which is 16.6%, the cutoff of importance.

**Supplemental Tables**

**Table S1** List of strains used in this study

| **Strain** | **Description** | **Source** |
| --- | --- | --- |
| DH5α | F-, 80dlacZM15, (*lacZYA*-*argF*)U169, *deoR*, *recA*1 *endA*1, *hsdR*17(rK- mK+), *phoA*, *supE*44, -, *thi*-1, *g*yrA96, *rel*A1 |  |
| MG1655(DE3) | F-, lamda-, ilvG-, rfb-50, rph-1 |  |
| *Ds*Red.GFPuv:2 plasmid system | MG1655(DE3) carrying pBluescript_P_T7_-*Ds*Red and pCDF2_P_tac_-GFPuv | This study |
| *Ds*Red.GFPuv:1 plasmid system, 200 copy | MG1655(DE3) carrying pBS_P_T7_-*Ds*Red.GFPuv | This study |
| *Ds*Red.GFPuv:1 plasmid system, 15-20 copy | MG1655(DE3) carrying pTargetF_P_T7_-*Ds*Red.GFPuv | This study |
| *Ds*Red.GFPuv: genome integrated | MG1655(DE3) with *DsRed* and *GFPuv* under a constitutive T7 promoter, integrated in the genome at the *araA* locus. | This study |
| WT | MG1655(DE3) carrying the pCDF2_P_tac_*-ispA(S80F).tObGES* | This study |
| *araA*::P_103_-*SIDF*/*pykF*::P_103_-*RGHE* | MG1655(DE3) with the genome-integrated *araA*::*SIDF* under the K1585103 promoter and *pykF::RGHE* under the K1585103 promoter. The strain also carries the pCDF2_P_tac_-*ispA(S80F).tObGES*. | This study |
| *araA*::P_103_-*SIDF*/*pykF*::P_105_-*RGHE* | MG1655(DE3) with the genome-integrated *araA*::*SIDF* under the K1585103 promoter and *pykF::RGHE* under the K1585105 promoter. The strain also carries the pCDF2_P_tac_-*ispA(S80F).tObGES*. | This study |
| *araA*::P_103_-*SIDF*/*pykF*::P_106_-*RGHE* | MG1655(DE3) with the genome-integrated *araA*::*SIDF* under the K1585103 promoter and *pykF::RGHE* under the K1585106 promoter. The strain also carries the pCDF2_P_tac_-*ispA(S80F).tObGES*. | This study |
| *araA*::P_103_-*SIDF*/*pykF*::P_101_-*RGHE* | MG1655(DE3) with the genome-integrated *araA*::*SIDF* under the K1585103 promoter and *pykF::RGHE* under the K1585101 promoter. The strain also carries the pCDF2_P_tac_-*ispA(S80F).tObGES*. | This study |
| *araA*::P_103_-*SIDF*/*pykF*::P_100_-*RGHE* | MG1655(DE3) with the genome-integrated *araA*::*SIDF* under the K1585103 promoter and *pykF::RGHE* under the K1585100 promoter. The strain also carries the pCDF2_P_tac_-*ispA(S80F).tObGES*. | This study |
| *araA*::P_105_-*SIDF*/*pykF*::P_103_-*RGHE* | MG1655(DE3) with the genome-integrated *araA*::*SIDF* under the K1585105 promoter and *pykF::RGHE* under the K1585103 promoter. The strain also carries the pCDF2_P_tac_-*ispA(S80F).tObGES*. | This study |
| *araA*::P_105_-*SIDF*/*pykF*::P_105_-RGHE | MG1655(DE3) with the genome-integrated *araA*::*SIDF* under the K1585105 promoter and *pykF::RGHE* under the K1585105 promoter. The strain also carries the pCDF2_P_tac_-*ispA(S80F).tObGES*. | This study |
| *araA*::P_105_-SIDF/*pykF*::P_106_-*RGHE* | MG1655(DE3) with the genome-integrated *araA*::*SIDF* under the K1585105 promoter and *pykF::RGHE* under the K1585106 promoter. The strain also carries the pCDF2_P_tac_-*ispA(S80F).tObGES*. | This study |
| *araA*::P_105_-*SIDF*/*pykF*::P_101_-*RGHE* | MG1655(DE3) with the genome-integrated *araA*::*SIDF* under the K1585105 promoter and *pykF::RGHE* under the K1585101 promoter. The strain also carries the pCDF2_P_tac_-*ispA(S80F).tObGES*. | This study |
| *araA*::P_105_-*SIDF*/*pykF*::P_100_-*RGHE* | MG1655(DE3) with the genome-integrated *araA*::*SIDF* under the K1585105 promoter and *pykF::RGHE* under the K1585100 promoter. The strain also carries the pCDF2_P_tac_-*ispA(S80F).tObGES*. | This study |
| *araA*::P_106_-*SIDF*/*pykF*::P_103_-*RGHE* | MG1655(DE3) with the genome-integrated *araA*::*SIDF* under the K1585106 promoter and *pykF::RGHE* under the K1585103 promoter. The strain also carries the pCDF2_P_tac_-*ispA(S80F).tObGES*. | This study |
| *araA*::P_106_-*SIDF*/*pykF*::P_105_-*RGHE* | MG1655(DE3) with the genome-integrated *araA*::*SIDF* under the K1585106 promoter and *pykF::RGHE* under the K1585105 promoter. The strain also carries the pCDF2_P_tac_-*ispA(S80F).tObGES*. | This study |
| *araA*::P_106_-*SIDF*/*pykF*::P_106_-*RGHE* | MG1655(DE3) with the genome-integrated *araA*::*SIDF* under the K1585106 promoter and *pykF::RGHE* under the K1585106 promoter. The strain also carries the pCDF2_P_tac_-*ispA(S80F).tObGES*. | This study |
| *araA*::P_106_-*SIDF*/*pykF*::P_101_-*RGHE* | MG1655(DE3) with the genome-integrated *araA*::*SIDF* under the K1585106 promoter and *pykF::RGHE* under the K1585101 promoter. The strain also carries the pCDF2_P_tac_-*ispA(S80F).tObGES*. | This study |
| *araA*::P_106_-*SIDF*/*pykF*::P_100_-*RGHE* | MG1655(DE3) with the genome-integrated *araA*::*SIDF* under the K1585106 promoter and *pykF::RGHE* under the K1585100 promoter. The strain also carries the pCDF2_P_tac_-*ispA(S80F).tObGES*. | This study |
| *araA*::P_106_-*SIDF*/*pykF*::P_tac_-*RGHE* | MG1655(DE3) with the genome-integrated *araA*::*SIDF* under the K1585106 promoter and *pykF::RGHE* under the P_tac_ promoter. The strain also carries the pCDF2_P_tac_-*ispA(S80F).tObGES*. | This study |
| *araA*::P_106_-*SIDF*/*pykF*::P_T7-lacO_-*RGHE* | MG1655(DE3) with the genome-integrated *araA*::*SIDF* under the K1585106 promoter and *pykF::RGHE* under the IPTG inducible T7-lacO promoter. The strain also carries the pCDF2_P_tac_-*ispA(S80F).tObGES*. | This study |
| *araA*::P_101_-*SIDF*/*pykF*::P_103_-*RGHE* | MG1655(DE3) with the genome-integrated *araA*::*SIDF* under the K1585101 promoter and *pykF::RGHE* under the K1585103 promoter. The strain also carries the pCDF2_P_tac_-*ispA(S80F).tObGES*. | This study |
| *araA*::P_101_-*SIDF*/*pykF*::P_105_-*RGHE* | MG1655(DE3) with the genome-integrated *araA*::*SIDF* under the K1585101 promoter and *pykF::RGHE* under the K1585105 promoter. The strain also carries the pCDF2_P_tac_-*ispA(S80F).tObGES*. | This study |
| *araA*::P_101_-*SIDF*/*pykF*::P_106_-*RGHE* | MG1655(DE3) with the genome-integrated *araA*::*SIDF* under the K1585101 promoter and *pykF::RGHE* under the K1585106 promoter. The strain also carries the pCDF2_P_tac_-*ispA(S80F).tObGES*. | This study |
| *araA*::P_101_-*SIDF*/*pykF*::P_101_-*RGHE* | MG1655(DE3) with the genome-integrated *araA*::*SIDF* under the K1585101 promoter and *pykF::RGHE* under the K1585101 promoter. The strain also carries the pCDF2_P_tac_-*ispA(S80F).tObGES*. | This study |
| *araA*::P_101_-*SIDF*/*pykF*::P_100_-*RGHE* | MG1655(DE3) with the genome-integrated *araA*::*SIDF* under the K1585101 promoter and *pykF::RGHE* under the K1585100 promoter. The strain also carries the pCDF2_P_tac_-*ispA(S80F).tObGES*. | This study |
| *araA*::P_100_-*SIDF*/*pykF*::P_103_-*RGHE* | MG1655(DE3) with the genome-integrated *araA*::*SIDF* under the K1585100 promoter and *pykF::RGHE* under the K1585103 promoter. The strain also carries the pCDF2_P_tac_-*ispA(S80F).tObGES*. | This study |
| *araA*::P_100_-*SIDF*/*pykF*::P_105_-*RGHE* | MG1655(DE3) with the genome-integrated *araA*::*SIDF* under the K1585100 promoter and *pykF::RGHE* under the K1585105 promoter. The strain also carries the pCDF2_P_tac_-*ispA(S80F).tObGES*. | This study |
| *araA*::P_100_-*SIDF*/*pykF*::P_106_-*RGHE* | MG1655(DE3) with the genome-integrated *araA*::*SIDF* under the K1585100 promoter and *pykF::RGHE* under the K1585106 promoter. The strain also carries the pCDF2_P_tac_-*ispA(S80F).tObGES*. | This study |
| *araA*::P_100_-*SIDF*/*pykF*::P_101_-*RGHE* | MG1655(DE3) with the genome-integrated *araA*::*SIDF* under the K1585100 promoter and *pykF::RGHE* under the K1585101 promoter. The strain also carries the pCDF2_P_tac_-*ispA(S80F).tObGES*. | This study |
| *araA*::P_100_-*SIDF*/*pykF*::P_100_-*RGHE* | MG1655(DE3) with the genome-integrated *araA*::*SIDF* under the K1585100 promoter and *pykF::RGHE* under the K1585100 promoter. The strain also carries the pCDF2_P_tac_-*ispA(S80F).tObGES*. | This study |
| *araA*::P_106_-*SIDF*/*pykF*::P_tac_-*RGHE-v2* | MG1655(DE3) with the genome-integrated *araA*::*SIDF* under the K1585106 promoter and *pykF::RGHE* under the P_tac_ promoter. The strain also carries the pCDFDuet1-kan^R^_P_T7-lacO_-*ispA(S80F)*.*tObGES* | This study |
| Geraniol:3 plasmids | MG1655(DE3) transformed with plasmids pIB_*araA::P_106_-SIDF,* pTargetF_*pykF::P_tac_-RGHE,* and pCDFDuet1-kan^R^_P_T7-lacO_-*ispA(S80F)*.*tObGES* | This study |

**Table S2** List of plasmids

| **Plasmid** | **Description** | **Source** |
| --- | --- | --- |
| pBluescript KS II+ | ColE1, *bla*, P_T7_ | Agilent |
| pCDFDuet1 | CloDF13, *aadA*, *lacI^q^*, P_T7-lacO_ | EMD Millipore |
| pCDFDuet1-kan^R^ | CloDF13, *kan^R^*, *lacI^q^*, P_T7-lacO_ | This study |
| pCDF2 | CloDF13, *aadA*, *lacI^q^, P_tac_* | EMD Millipore |
| pKD46_*Cas9.RecA.Cure* | *SpCas9, PaRecA, repA101ts, bla, araC*, *araB*p-λ γ -λ β -λex | Gift from Dr. Quanjiang Ji |
| pYTK001 | Rep_ori, *cml*, *sfGFP* dropout | (Lee et al. 2015) |
| pTargetF | pMB1, *aadA*, P_J23119_ | (Jiang et al. 2015) |
| pTargetF_*araA* | pTargetF containing a sgRNA to target the *araA* locus | This study |
| pTargetF_*pykF* | pTargetF containing a sgRNA to target the *pykF* locus | This study |
| pBluescript KS II+_P_T7_-*DsRed* | pBluescript KS II+ expressing *DsRed* from a constitutive T7 promoter. | This study |
| pCDF2_P_tac_-*GFPuv* | Plasmid pCDF2 expressing *GFPuv* from a P_tac_ promoter. | This study |
| pBluescript KS II+_P_T7_-*DsRed.GFPuv* | pBluescript KS II+ expressing *DsRed* and *GFPuv* as an operon from a constitutive T7 promoter | This study |
| pTargetF_*araA::*P_T7_-*DsRed.GFPuv* | pTargetF_*araA* modified to express *DsRed* and *GFPuv* as an operon from a constitutive T7 promoter and the homology regions of the *araA* locus | This study |
| pCDF2_P_tac_-*ispA(S80F).tObGES* | Plasmid pCDF2 expressing *ispA(S80F)* and *tObGES* from a double P_tac_ promoter | This study |
| pTargetF_*araA GG* | Plasmid pTargetF_*araA* modified to include two BsaI sites | This study |
| pTargetF_*pykF GG* | Plasmid pTargetF_*pykF* modified to include two BsaI sites | This study |
| pYTK001_*sfGFP dropout* | Plasmid pYTK001 expressing a *sfGFP* transcription unit for assembling pTargetF_*araA/pykF int GG1* | This study |
| pYTK001_*Kan^R^ dropout* | Plasmid pYTK001 contain a kanamycin resistant gene (*kan^r^*) for assembling pTargetF_*araA/pykF int GG2* | This study |
| pYTK001_*sfGFP dropout2* | Plasmid pYTK001 modified to contain *sfGFP* with different overhangs than pYTK001_*sfGFP dropout* for assembling pTargetF_*araA/pykF int GG3* | This study |
| pYTK001_*LR-araA* | pYTK001 containing the 5’ homology region of the *araA* locus, flanked by two BsaI sites | This study |
| pYTK001_*RR-araA* | pYTK001 containing the 3’ homology region of the *araA* locus, flanked by two BsaI sites | This study |
| pYTK001_*dxs* | pYTK001 containing the *dxs* flanked by two BsaI sites | This study |
| pYTK001_*idi* | pYTK001 containing *idi* flanked by two BsaI sites | This study |
| pYTK001_*ispDF-term* | pYTK001 containing *ispDF* linked to L3S2P21 terminator sequence, flanked by two BsaI sites | This study |
| pYTK001_*NAT* | pYTK001 containing a nourseothricin resistant gene (*NAT^R^*) transcription unit flanked by two BsaI sites. | This study |
| pYTK001_*LR-pykF* | pYTK001 containing the 5’ homology region of the *pykF* locus, flanked by two BsaI sites | This study |
| pYTK001_*RR-pykF* | pYTK001 containing the 3’ homology region of the *pykF* locus, flanked by two BsaI sites | This study |
| pYTK001_*dxr* | pYTK001 containing the *dxr* flanked by two BsaI sites | This study |
| pYTK001_*ispG* | pYTK001 containing the *ispG* flanked by two BsaI sites | This study |
| pYTK001_*ispH* | pYTK001 containing the *ispH* flanked by two BsaI sites | This study |
| pYTK001_*ispE-term* | pYTK001 containing the *ispE* linked to T_PheA_ terminator, flanked by two BsaI sites | This study |
| pYTK001_P_T7_-*lacI* | pYTK001 containing *lacI* under a constitutive T7 promoter, flanked by two BsaI sites | This study |
| pYTK001_P_tac_ | pYTK001 containing an IPTG-inducible P_tac_ promoter, flanked by two BsaI sites | This study |
| pTargetF_*araA int GG1* | pTargetF_*araA GG* modified to contain the 5’ homologous region for the *araA* locus, *sfGFP*, and the 3’ homologous region for the *araA* locus. BsaI sites are at the 5’ end of the 5’ homology region and the 3’ end of the 3’ homology region. | This study |
| pTargetF_*araA int GG2* | pTargetF_*araA int GG1* with the *sfGFP* replaced by a *kan^R^*, *ispDF* with its RBS and a terminator, and the nourseothricin resistant gene (*NAT^R^*). BsaI sites are at the 5’ end of the *kan^R^* and 3’ end of the *NAT^R^*. | This study |
| pTargetF_*araA int GG3* | pTargetF_*araA int GG2* with the *kan^R^* replaced by *sfGFP*, *dxs* with its RBS, and *idi* with its RBS. BsaI sites are at the 5’ end of the *sfGFP* and 3’ end of *idi*. | This study |
| pTargetF_*pykF int GG1* | pTargetF_*pykF GG* modified to encode the 5’ homologous region for the *pykF* locus, *sfGFP*, and the 3’ homology region for the *pykF* locus. BsaI sites are at the 5’ end of the 5’ homology region and the 3’ end of the 3’ homologous region. | This study |
| pTargetF_*pykF int GG2* | pTargetF_*pykF int GG1* with the *sfGFP* replaced by the *kan^R^*, *ispH* with its RBS, and *ispE* with its RBS and terminator. BsaI sites are at the 5’ end of the *kan^R^* and 3’ end of the terminator. | This study |
| pTargetF_*pykF int GG3* | pTargetF_*pykF int GG2* with the *kan^R^* replaced by *sfGFP*, *dxr* with its RBS, and *ispG* with its RBS. BsaI sites are at the 5’ end of the *sfGFP* and 3’ end of *ispG*. | This study |
| pTargetF_ *araA::P_103_-SIDF* | pTargetF_*araA int GG3* with the *sfGFP* replaced by *lacI* under a constitutive T7 promoter, and the K1585103 promoter. | This study |
| pTargetF_ *araA::P_105_-SIDF* | pTargetF_*araA int GG3* with the *sfGFP* replaced by *lacI* under constitutive T7 promoter, and the K1585105 promoter. | This study |
| pTargetF_ *araA::P_106_-SIDF* | pTargetF_*araA int GG3* with the *sfGFP* replaced by *lacI* under constitutive T7 promoter, and the K1585106 promoter. | This study |
| pTargetF_ *araA::P_101_-SIDF* | pTargetF_*araA int GG3* with the *sfGFP* replaced by *lacI* under constitutive T7 promoter, and the K1585101 promoter. | This study |
| pTargetF_ *araA::P_100_-SIDF* | pTargetF_*araA int GG3* with the *sfGFP* replaced by *lacI* under constitutive T7 promoter, and the K1585100 promoter. | This study |
| pTargetF_*pykF::P_103_-RGHE* | pTargetF_*pykF int GG3* with the *sfGFP* replaced by *lacI* under constitutive T7 promoter, and the K1585103 promoter. | This study |
| pTargetF_ *pykF::P_105_-RGHE* | pTargetF_*pykF int GG3* with the *sfGFP* replaced by *lacI* under constitutive T7 promoter, and the K1585105 promoter. | This study |
| pTargetF_ *pykF::P_106_-RGHE* | pTargetF_*pykF int GG3* with the *sfGFP* replaced by *lacI* under constitutive T7 promoter, and the K1585106 promoter. | This study |
| pTargetF_ *pykF::P_101_-RGHE* | pTargetF_*pykF int GG3* with the *sfGFP* replaced by *lacI* under constitutive T7 promoter, and the K1585101 promoter. | This study |
| pTargetF_ *pykF::P_100_-RGHE* | pTargetF_*pykF int GG3* with the *sfGFP* replaced by *lacI* under constitutive T7 promoter, and the K1585100 promoter. | This study |
| pTargetF_ *pykF::P_tac_-RGHE* | pTargetF_*pykF int GG3* with the *sfGFP* replaced by *lacI* under constitutive T7 promoter, and the P_tac_ promoter. | This study |
| pTargetF_ *pykF::P_T7-lacO_-RGHE* | pTargetF_*pykF int GG3* with the *sfGFP* replaced by *lacI* under constitutive T7 promoter, and the IPTG-inducible T7-lacO promoter. | This study |
| pIB_ *araA::P_106_-SIDF* | p15A, *cml*, *araA*::SIDF under the K1585106 promoter. | This study |
| pCDFDuet1-kan^R^_P_T7-lacO_-*ispA(S80F)*.*tObGES* | Plasmid pCDFDuet1 modified to contain *kan^R^*, and *ispA(S80F)* and *tObGES* under a T7-lacO promoter | This study |

**Table S3** List of primers for cloning

RBS sequences for all genes except *ispG*, *ispA(S80F)*, and t*Ob*GES are highlighted in the primer sequences.

RBS sequence for *ispG*: gccagccaataaggagatttcact

RBS sequence for *ispA(S80F)*: taagtatacaaaaattttaaagataaggaggtaaagt

RBS sequence for t*Ob*GES: cacaagcaataaggagcgattccat

| Primer | Sequence 5’ 🡪 3’ (RBS sequences highlighted) | Notes |
| --- | --- | --- |
| *Ds*Red_RBS FW | tcggttaggagaagcagcccgcgctgagaaataaggaggttttttatggacaacaccgaggac | For amplifying *DsRed* to clone pBluescript KS II+_P_T7_-*DsRed* |
| *Ds*Red RW | tcactcgagctgggagcc | For amplifying *DsRed* to clone pBluescript KS II+_P_T7_-*DsRed* cloning |
| pBS_*Ds*Red FW | ccggctcccagctcgagtgacgaattggagctccacc | For amplifying the plasmid backbone to clone pBluescript KS II+_ P_T7_-*DsRed* cloning |
| pBS_*Ds*Red RW | gggctgcttctcctaaccgaccctatagtgagtcgtattacg | For amplifying the plasmid backbone to clone pBluescript KS II+_ P_T7_-*DsRed* cloning |
| GFPuv_RBS FW | tggcagttgaactggatctgagatttaaggaggtattttatgagtaaaggagaagaac | For amplifying *GFPuv* to clone pCDF2_P_tac_-*GFPuv* |
| GFPuv RW | ttatttgtagagctcatcc | For amplifying *GFPuv* to clone pCDF2_P_tac_-*GFPuv* |
| pCDF2_GFPuv FW | tggatgagctctacaaataattatttgccgactaccttg | For amplifying the pCDF2 backbone to clone pCDF2_P_tac_-*GFPuv* |
| pCDF2_GFPuv RW | cagatccagttcaactgccagatcctgtttcctgtgtg | For amplifying the pCDF2 backbone to clone pCDF2_P_tac_-*GFPuv* |
| pBS_GFPuv FW | tggatgagctctacaaataacgaattggagctccacc | For cloning pBluescript KS II+_P_T7_-*DsRed.GFPuv* |
| dsRed_GFPuv RW | cagatccagttcaactgccatcactcgagctgggagcc | For cloning pBluescript KS II+_P_T7_-*DsRed.GFPuv* |
| LR-araA FW | agtgcacgcagacatcc | For cloning pTargetF_*araA::*P_T7_-*DsRed.GFPuv* |
| LR-araA_RR RW | cgatactgtcccacggcagcttacggtttgttgagcatggt | For cloning pTargetF_*araA::*P_T7_-*DsRed.GFPuv* |
| RR-araA FW | gctgccgtgggacagtatcgatat | For cloning pTargetF_*araA::*P_T7_-*DsRed.GFPuv* |
| RR-araA RW | aggtggtggtgaacgcgtg | For cloning pTargetF_*araA::*P_T7_-*DsRed.GFPuv* |
| pTargetF_RR FW | ccacgcgttcaccaccacctatctattaccctgttatccc | For cloning pTargetF_*araA::*P_T7_-*DsRed.GFPuv* |
| pTargetF_LR RW | atgggatgtctgcgtgcactctaagcttctgcaggt | For cloning pTargetF_*araA::*P_T7_-*DsRed.GFPuv* |
| RR_pTar_GFPuv FW | tggatgagctctacaaataagctgccgtgggacagtatcgatat | For cloning pTargetF_*araA::*P_T7_-*DsRed.GFPuv* |
| LR_pTar_*Ds*Red RW | cctatagtgagtcgtattacttacggtttgttgagcatggtcagg | For cloning pTargetF_*araA::*P_T7_-*DsRed.GFPuv* |
| pTargetF-sgRNA FW | P-attccacacccagttcaacggttttagagctagaaatagc | For cloning pTargetF_*araA* |
| pTar-pykF sgRNA FW(P) | P-gagcacctgaaagcgcacgggttttagagctagaaatagc | For cloning pTargetF_*pykF* |
| pTargetF GG FW (P) | P-tttgacggtctctccgaattaccctg | For cloning pTargetF_*araA GG* and pTargetF_*pykF GG* |
| pTargetF GG RW (P) | P-tttgatggtctctactcctaagcttctg | For cloning pTargetF_*araA GG* and pTargetF_*pykF GG* |
| pYTK001 FW GG | tttcgtctctgaccagaccaataaaaaacgccc | For amplifying the pYTK001 backbone to clone pYTK001_*sfGFP dropout* |
| pYTK001 RW GG | tttcgtctcgccgactacggttatccacagaatc | For amplifying the pYTK001 backbone to clone pYTK001_*sfGFP dropout* |
| sfGFP FW GG | tttcgtctcgtcggccctagagaccgacggaaagtgaaacgtgatttcat | For cloning pYTK001_*sfGFP dropout* |
| sfGFP RW GG | tttcgtctctggtctgtatgagaccgacgtataaacgcagaaaggcc | For cloning pYTK001_sfGFP dropou*t* |
| KanR FW GG | tttcgtctcgtcggccctagagacccaccgacgtcggaattgc | For cloning pYTK001_*Kan^R^ dropout* |
| KanR RW GG | tttcgtctctggtcggattgagacctgactagtgcttggattctcacc | For cloning pYTK001_ *Kan^R^ dropout* |
| sfGFP RW GG2 | tttcgtctctggtcgccatgagaccgacgtataaacgcagaaaggcc | For cloning pYTK001_sfGFP dropout2 |
| araA-LR FW | tttcgtctcgtcggggtctcggagtagtgcacgcagacat | For cloning pYTK001_*LR-araA* |
| araA-LR RW | tttcgtctctggtcggtctctagggttacggtttgttgagcatg | For cloning pYTK001_*LR-araA* |
| RR-araA FW GG | tttcgtctcgtcggggtctcgtacagctgccgtgggacagta | For cloning pYTK001_*RR-araA* |
| RR-araA mut RW | tttcgtctcccgcctgacgcatcca | For domesticating *araA* type II restriction sites to clone pYTK001_*RR-araA* |
| RR-araA mut FW | tttcgtctcaggcggtatctaaacaggata | For domesticating *araA* type II restriction sites to clone pYTK001_*RR-araA* |
| RR-araA RW GG | tttcgtctctggtcggtctcttcggaggtggtggtgaacgc | For cloning pYTK001_*RR-araA* |
| dxs FW GG | tttcgtctcgtcggggtctcatggcgagcggacaaccgaatttagttaaggagcactttatgagttttgatattgccaaataccc | For cloning pYTK001_*dxs* |
| dxs RW GG | tttcgtctctggtcggtctctcgttttatgccagccaggcct | For cloning pYTK001_*dxs* |
| idi FW GG | tttcgtctcgtcggggtctcgaacgggaataatagtaaggaaaggtaatttaatgcaaacggaacacgt | For cloning pYTK001_*idi* |
| idi RW GG | tttcgtctctggtcggtctctggatttatttaagctgggtaaatgcag | For cloning pYTK001_*idi* |
| ispDF FW GG | tttcgtctcgtcggggtctcgatccacgagctagttaataatatataaggaggtatttgatggcaaccactcatttgga | For cloning pYTK001_*ispDF-term* |
| ispDF-term RW GG | tttcgtctctggtcggtctctcagcggaccaaaacgaaaaaaggcccccctttcgggaggcctcttttctggaatttggtaccgagtcattttgttgccttaatgagtagc | For cloning pYTK001_*ispDF-term* |
| NAT FW GG | tttcgtctcgtcggggtctcggctgtaggtctagagatctgtttagcttg | For cloning pYTK001_*NAT* |
| NAT mut RW | tttcgtctcgagtacgatacgaccacga | For domesticating *NAT* type II restriction sites to clone pYTK001_*NAT* |
| NAT mut FW | tttcgtctcgtactccggctg | For domesticating *NAT* type II restriction sites to clone pYTK001_*NAT* |
| NAT RW GG | tttcgtctctggtcggtctcttgtaattaagggttctcgagagctc | For cloning pYTK001_*NAT* |
| pykF-LR FW | gattctgtggataaccgtagggtctcggagtgaacttctctcatggtgactatgc | For cloning pYTK001_*LR-pykF* |
| pykF-LR RW | cgttttttattggtctggtcggtctctagggttagatttcgataacgtcagaacgc | For cloning pYTK001_*LR-pykF* |
| pykF-RR FW | gattctgtggataaccgtagggtctcgtacagaaaacatccacatcatctcc | For cloning pYTK001_*RR-pykF* |
| pykF-RR RW | cgttttttattggtctggtcggtctcttcggatttaccgccctgagtag | For cloning pYTK001_*RR-pykF* |
| dxr FW GG2 | tttcgtctcgtcggggtctcatatgagctacttggataacatacggaggaaaactatgaagcaactcaccattctgg | For cloning pYTK001_*dxr* |
| dxr mut RW | tttcgtctccaatggcgtttcacggaaag | For domesticating *dxr* type II restriction sites to clone pYTK001_*dxr* |
| dxr mut FW | tttcgtctcccattgcgcgatttggcaacaatgacg | For domesticating *dxr* type II restriction sites to clone pYTK001_*dxr* |
| dxr RW GG | tttcgtctctggtcggtctctcgtttcagcttgcaagacgcat | For cloning pYTK001_*dxr* |
| ispG FW GG2 | tttcgtctcgtcggggtctcgaacggccagccaataaggagatttc | For cloning pYTK001_*ispG* |
| ispG RW GG2 | tttcgtctctggtcggtctctggatttatttttcaacctgctgaacgt | For cloning pYTK001_*ispG* |
| ispH FW | gattctgtggataaccgtagggtctcgatcctaagaaatcgactaaggacaatttacatgcagatcctgttggcc | For cloning pYTK001_*ispH* |
| ispH RW | cgttttttattggtctggtcggtctctcagcttaatcgacttcacgaatatcgac | For cloning pYTK001_*ispH* |
| ispE FW | tttcgtctcgtcggggtctcggctgcaataaaacgataagaagaggcaatttatgcggacacagtggccctctccg | For cloning pYTK001_*ispE-term* |
| ispE mut RW | tttcgtctcagaccgccgcccatcggcaaa | For domesticating *ispE* type II restriction sites to clone pYTK001_*ispE-term* |
| ispE mut FW | tttcgtctccggtctaggcggtggttcat | For domesticating *ispE* type II restriction sites to clone pYTK001_*ispE-term* |
| ispE-term RW | tttcgtctctggtcggtctcttgtattttgttatcaataaaaaaggccccccgatttgggaggccttattgttcgtcttaaagcatggctctgtgcaatgggg | For cloning pYTK001_*ispE-term* |
| LacI-T7 FW Gib | gattctgtggataaccgtagggtctcgccctgtaatacgactcactataggggaggtaaagaacccggaggaggtattttcgtgaaaccagtaacgttatacgatg | For cloning pYTK001_P_T7_-*lacI* |
| LacI RW Gib | cgttttttattggtctggtcggtctctcatagaaccgttatgatgtcggcg | For cloning pYTK001_P_T7_-*lacI* |
| tac FW GG 3 | tttcgtctcgtcggggtctcatatggagctgttgacaattaatcatcggctcg | For cloning pYTK001_P_tac_ |
| tac RW GG | tttcgtctctggtcggtctcagccagatcctgtttcctgtgtg | For cloning pYTK001_P_tac_ |
| T7 FW GG | tttcgtctcgtcggggtctcatatgttaatacgactcactataggggaattgtgagcggataacaattccctggctgagaccgaccagagacgaaa | Oligo for pYTK001_P_T7-lacO_. The *lacO* region is underlined. |
| T7 RW GG | tttcgtctctggtcggtctcagccagggaattgttatccgctcacaattcccctatagtgagtcgtattaacatatgagaccccgacgagacgaaa | Oligo for pYTK001_P_T7-lacO_ |
| K1585103 double-stranded oligo | tttcgtctcgtcggggtctcatatgctgatagctagctcagtcctagggattatgctagctactagagtgtgtggaattgtgagcggataacaatttcacacatggctgagaccgaccagagacgaaa | Oligo for K1585103 promoter. The *lacO* region is underlined. |
| K1585105 double-stranded oligo | tttcgtctcgtcggggtctcatatgtttacggctagctcagtcctaggtactatgctagctactagagtgtgtggaattgtgagcggataacaatttcacacatggctgagaccgaccagagacgaaa | Oligo for K1585105 promoter. The *lacO* region is underlined. |
| K1585106 double-stranded oligo | tttcgtctcgtcggggtctcatatgtttacggctagctcagtcctaggtatagtgctagctactagagtgtgtggaattgtgagcggataacaatttcacacatggctgagaccgaccagagacgaaa | Oligo for K1585106 promoter. The *lacO* region is underlined. |
| K1585101 double-stranded oligo | tttcgtctcgtcggggtctcatatgtttacagctagctcagtcctaggtattatgctagctactagagtgtgtggaattgtgagcggataacaatttcacacatggctgagaccgaccagagacgaaa | Oligo for K1585101 promoter. The *lacO* region is underlined. |
| K1585100 double-stranded oligo | tttcgtctcgtcggggtctcatatgttgacggctagctcagtcctaggtacagtgctagctactagagtgtgtggaattgtgagcggataacaatttcacacatggctgagaccgaccagagacgaaa | Oligo for K1585100 promoter. The *lacO* region is underlined. |
| p15A-Cm FW Gib | tattgagctggggatgaagccctcgctctgctaatcctgttac | For cloning pIB_*araA::P_106_-SIDF* |
| p15A-Cm RW Gib | caggtcgactctagagaattgatcgggctcgccacttc | For cloning pIB_*araA::P_106_-SIDF* |
| Km FW Gib | tcacactgcttccggtagtccaccgacgtcggaattg | For cloning pCDFDuet1-kan^R^ |
| Km RW Gib | ccgagtgagctagctatttgctagtgcttggattctcacc | For cloning pCDFDuet1-kan^R^ |
| pCDFDuet1 FW | caaatagctagctcactcgg | For cloning pCDFDuet1-kan^R^ |
| pCDFDuet1 RW | gactaccggaagcagtgt | For cloning pCDFDuet1-kan^R^ |
| ispA* FW Gib | caccatcatcaccacagccaggatcctaagtatacaaaaattttaaag | For cloning pCDFDuet1-kan^R^_P_T7-lacO_-*ispA(S80F)*.*tObGES* |
| *Ob*GES RW Gib | ggtttctttaccagactcgaaagcttttattgtgtaaaaaacagggc | For cloning pCDFDuet1-kan^R^_P_T7-lacO_-*ispA(S80F)*.*tObGES* |

**Table S4** List of primers for genotyping

| Primer | 5’ 🡪 3’ sequence | Purpose | Notes |
| --- | --- | --- | --- |
| gen CmR FW | aaccaggtcattatgcaggc | Forward primer to screen for the 5’ end of *araA*::SIDF | Binds to the genome region upstream of *araA::SIDF* |
| gen CmR RW | caggcgttacataccggatg | Reverse primer to screen for the 3’ end of *araA*::SIDF | Binds to the genome region downstream of *araA::SIDF* |
| dxs RW_short | ttatgccagccaggccttg | Reverse primer to screen for the 5’ end of *araA*::SIDF | Binds to the *dxs* coding region |
| pBS-araA seq P9 FW | gccatatggatgatgttaacgt | Forward primer to screen for the 3’ end of *araA*::SIDF | Binds to the *ispF* coding region |
| gen. pykF FW | gctggcatgaacgttatgc | Forward primer to screen for the 5’ end of the *pykF*::RGHE | Binds to the genome region upstream of *pykF::RGHE* |
| gen. pykF RW2 | gcagaacgttcagacaatgc | Reverse primer to screen for the 3’ end of the *pykF*::RGHE | Binds to the genome region downstream of *pykF::RGHE* |
| dxr RW_screen2 | tcgtccattacggcatagc | Reverse primer to screen for the 5’ end of the *pykF*::RGHE | Binds to the *dxr* coding region |
| ispE FW_screen2 | gatacagagtctgaagcccg | Forward primer to screen for the 3’ end of the *pykF*::RGHE | Binds to the *ispE* coding region |

**Table S5** sgRNA protospacer sequences

| Target locus | 5’ 🡪3’ sequence |
| --- | --- |
| *araA* | attccacacccagttcaacg |
| *pykF* | gagcacctgaaagcgcacgg |

**Table S6** Homology regions for integration of MEP genes at *araA* and *pykF* genomic loci

| Homology region | Sequence |
| --- | --- |
| 5’ homology region of *araA* locus | agtgcacgcagacatcccatcagctcagcaaaaaatggccagtgcggtagagaaaaccctgcaaccgtgcagcgagcaggcacaacgctttgaacagctttatcgccgctatcagcaatgggcgatgagcgccgaacaacactatcttccaacttccgccccggcacaggctgcccaggccgttgcgactctataaggacacgataatgacgatttttgataattatgaagtgtggtttgtcattggcagccagcatctgtatggcccggaaaccctgcgtcaggtcacccaacatgccgagcacgtcgttaatgcgctgaatacggaagcgaaactgccctgcaaactggtgttgaaaccgctgggcaccacgccggatgaaatcaccgctatttgccgcgacgcgaattacgacgatcgttgcgctggtctggtggtgtggctgcacaccttctccccggccaaaatgtggatcaacggcctgaccatgctcaacaaaccgt |
| 3’ homology region of *araA* locus | gctgccgtgggacagtatcgatatggactttatgaacctgaaccagactgcacatggcggtcgcgagttcggcttcattggcgcgcgtatgcgtcagcaacatgccgtggttaccggtcactggcaggataaacaagcccatgagcgtatcggctcctggatgcgtcaggcggtctctaaacaggatacccgtcatctgaaagtctgccgatttggcgataacatgcgtgaagtggcggtcaccgatggcgataaagttgccgcacagatcaagttcggtttctccgtcaatacctgggcggttggcgatctggtgcaggtggtgaactccatcagcgacggcgatgttaacgcgctggtcgatgagtacgaaagctgctacaccatgacgcctgccacacaaatccacggcaaaaaacgacagaacgtgctggaagcggcgcgtattgagctggggatgaagcgtttcctggaacaaggtggcttccacgcgttcaccaccacct |
| 5’ homology region of *pykF* locus | gaacttctctcatggtgactatgcagaacacggtcagcgcattcagaatctgcgcaacgtgatgagcaaaactggtaaaaccgccgctatcctgcttgataccaaaggtccggaaatccgcaccatgaaactggaaggcggtaacgacgtttctctgaaagctggtcagacctttactttcaccactgataaatctgttatcggcaacagcgaaatggttgcggtaacgtatgaaggtttcactactgacctgtctgttggcaacaccgtactggttgacgatggtctgatcggtatggaagttaccgccattgaaggtaacaaagttatctgtaaagtgctgaacaacggtgacctgggcgaaaacaaaggtgtgaacctgcctggcgtttccattgctctgccagcactggctgaaaaagacaaacaggacctgatctttggttgcgaacaaggcgtagactttgttgctgcttcctttattcgtaagcgttctgacgttatcgaaatc |
| 3’ homology region of *pykF* locus | gaaaacatccacatcatctccaaaatcgaaaaccaggaaggcctcaacaacttcgacgaaatcctcgaagcctctgacggcatcatggttgcgcgtggcgacctgggtgtagaaatcccggtagaagaagttatcttcgcccagaagatgatgatcgaaaaatgtatccgtgcacgtaaagtcgttatcactgcgacccagatgctggattccatgatcaaaaacccacgcccgactcgcgcagaagccggtgacgttgcaaacgccatcctcgacggtactgacgcagtgatgctgtctggtgaatccgcaaaaggtaaatacccgctggaagcggtttctatcatggcgaccatctgcgaacgtaccgaccgcgtgatgaacagccgtctcgagttcaacaatgacaaccgtaaactgcgcattaccgaagcggtatgccgtggtgccgttgaaactgctgaaaaactggatgctccgctgatcgtggttgctactcagggcggtaaat |

**Supplemental codes**

**R code for principal component analysis (PCA)**

> library("FactoMineR")

> install.packages("devtools")

> library("devtools")

> install_github("kassambara/factoextra")

> library("factoextra")

> metabolites_data <- read.csv("D:\\Asus laptop complete backup\\D drive_New Volume\\Wang lab\\MEP project data\\MEP intermediates LCMS\\20221025 Intermediate quantification\\Integrated again and quantified\\metabolites.csv")	#reads the .csv file

> new_metabolites_dataset <- metabolites_data[,-1]	#reads the file without the first column

> rownames(new_metabolites_dataset) <- metabolites_data[,1]	#reads only the first column to assign “row names”

> View(new_metabolites_dataset)	#displays how the file is being read

> pca <- PCA(new_metabolites_dataset, graph = FALSE)	#correlation-based principal component analysis with scaling to unit variance

> fviz_pca_ind(pca, repel = TRUE)	#displays biplot

> fviz_eig(pca, addlabels = TRUE)	#displays scree plot

> var <- get_pca_var(pca)	#extracts the results for individual parameters of PCA

> a<-fviz_contrib(pca, choice = "var", axes = 1)	#assigns PC1 individual contributors graph to a variable

> b<-fviz_contrib(pca, choice = "var", axes = 2)	#assigns PC2 individual contributors graph to a variable

> install.packages("gridExtra")

> library(gridExtra)

> grid.arrange(a,b, ncol=2)	#displays PC1 and PC2’s individual contributors

**Appendix I**

4-nt overhangs used for Golden Gate assembly of MEP pathway genes:

ccct, tggc, aacg, atcc, gctg, taca, ccga

These overhangs are from left to right in the cloned plasmids.

**References:**

Jiang Y, Chen B, Duan C, Sun B, Yang J, Yang S (2015) Multigene editing in the *Escherichia coli* genome via the CRISPR-Cas9 system. Appl Environ Microbiol 81(7):2506-2514 doi:10.1128/AEM.04023-14

Lee ME, DeLoache WC, Cervantes B, Dueber JE (2015) A highly characterized yeast toolkit for modular, multipart assembly. ACS Synth Biol 4(9):975-986 doi:10.1021/sb500366v
